# Identification of regions required for allelic specificity at the cell wall remodeling allorecognition checkpoint in *Neurospora crassa*

**DOI:** 10.1101/2025.01.17.633681

**Authors:** Adriana M. Rico-Ramirez, N. Louise Glass

## Abstract

Allorecognition is the ability of organisms/cells to differentiate self from non-self. In the fungus *Neurospora crassa*, allorecognition systems serve as checkpoints to restrict germling/hyphal fusion between genetically incompatible strains. The <u>c</u>ell <u>w</u>all <u>r</u>emodeling (*cwr*) checkpoint functions after chemotrophic interactions and is triggered upon cell/hyphal contact, regulating cell wall dissolution and subsequent cell fusion. The *cwr* region consists of two linked loci, *cwr-1* and *cwr-2*, that are under severe linkage disequilibrium. Phylogenetic analysis of wild *N. crassa* populations showed that *cwr-1/cwr-2* alleles fall into six different haplogroups (HGs). Strains containing deletions of *cwr-1* and *cwr-2* will fuse with previously HG incompatible cells, indicating *cwr* negatively regulates cell fusion. CWR-1 encodes a polysaccharide monooxygenase (PMO) domain that oxidatively degrades chitin; the PMO domain is sufficient to cause fusion arrest and confers allelic specificity by interacting *in trans* with CWR-2, a predicted transmembrane protein. However, the catalytic activity of CWR-1 is not required for triggering a block in cell fusion. The L2 and LC regions of the CWR-1 PMO domain show high levels of structural variability between different HGs. CWR-1 chimeras containing a LC region from a different HG were sufficient to trigger a cell fusion block, but not quite at wild type levels, suggesting that the complete PMO structure is necessary for allorecognition. Modeling of the transmembrane protein CWR-2 revealed allelic variability in the two major extracellular domains (ED2/ED4). Chimeras of CWR-2 with swapped ED2 or ED4 or ED2/ED4 domains from different *cwr-2* haplogroups also altered allelic specificity.

**Summary:** Allorecognition or nonself recognition enables fungi to distinguish genetically different individuals, thereby regulating cooperation to form mycelial networks. This study focused on the cell wall remodeling checkpoint (*cwr*), where genetic differences in two genes, *cwr-1* and *cwr-2*, triggers allorecognition. Upon cell contact, CWR-1 in one cell functions *in trans* with CWR-2 in a second cell to confer a block in cell fusion. Chimeric proteins were created and tested to pinpoint domains involved in allelic specificity. For CWR-1, the L2 and LC domains are critical, while for CWR-2, the ED2 and ED4 domains have an important role in regulating cell fusion.

**Graphical Abstract.**
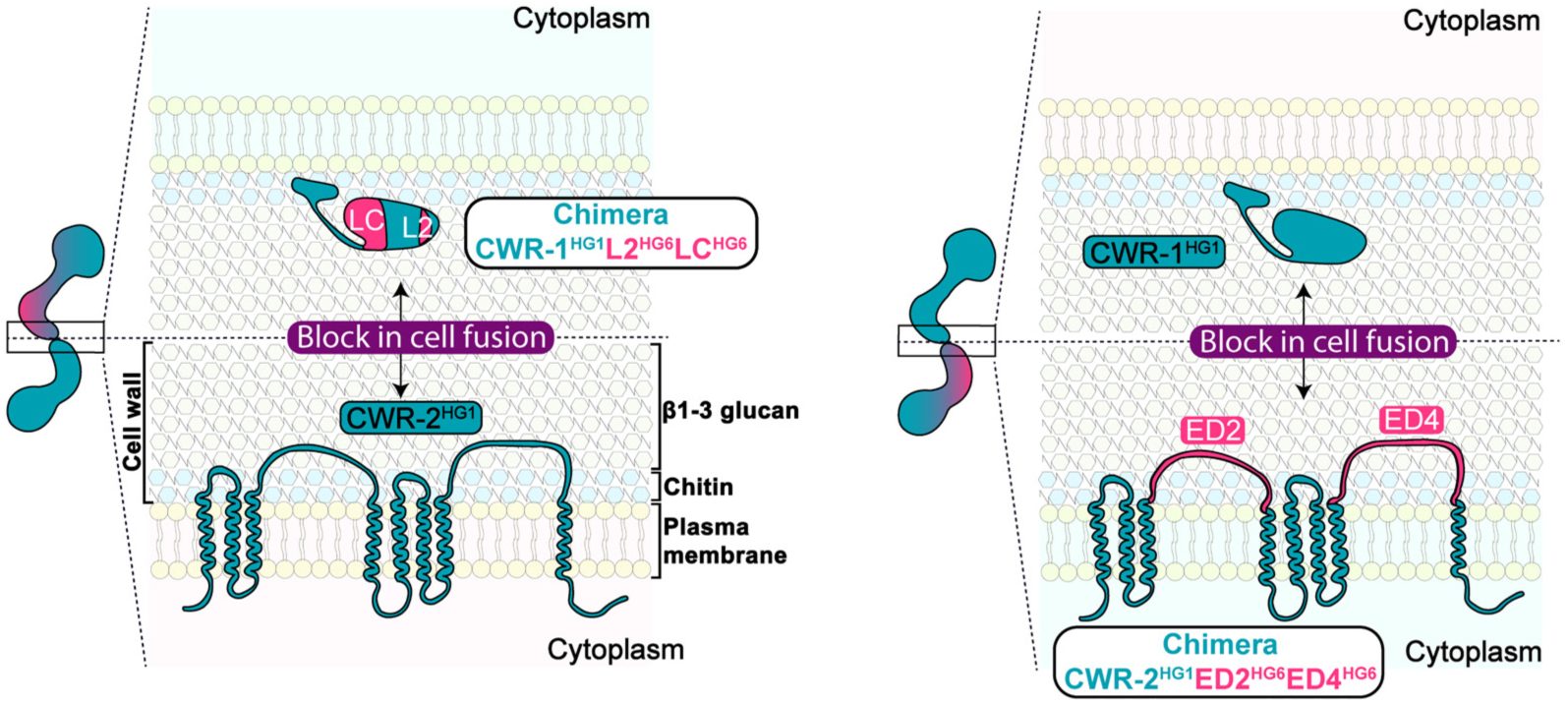

## Introduction

Allorecognition is defined as the capability of the organism to discern self from non-self. Allorecognition processes are highly specific, and their control depends on numerous genetic variables. In nature, there are many examples of species that use these systems, from vertebrates to bacteria. In vertebrates, the Major Histocompatibility Complex (MHC) is involved in nonself recognition, thereby triggering an immune response, particularly during organ and tissue transplantation (Afzali *et al*. 2008; Marino *et al*. 2016; Charmetant *et al*. 2024). The amoeba *Dictyostelium discoideum* uses allorecognition systems to restrict formation of aggregates between genetically different cells, which is required for the formation of multicellular sporulation structures under starvation conditions (Kessin 2001; Katoh-Kurasawa *et al*. 2024). In marine invertebrates, such as the colonial ascidian *Botryllus schlosseri* and cnidarian *Hydractinia symbiolongicarpus*, allorecognition restricts fusion of colonies to self and close kin, thus mediating the outcome of contested space in populations (Rosengarten and Nicotra 2011; Karadge *et al*. 2015; Huene *et al*. 2022; Rodriguez-Valbuena *et al*. 2024).

In filamentous fungi, allorecognition systems during vegetative growth are crucial for enabling fusion between cells (germlings and/or hyphae) that are genetically identical/similar, ensuring successful colony development and asexual reproduction. To achieve this, fungi have checkpoints that regulate the cell fusion process to avoid fusing with an unrelated individual that can potentially transfer detrimental elements, such as mycoviruses and debilitated organelles (Debets *et al*. 1994; Van Diepeningen *et al*. 1997; Debets and Griffiths 1998). In *Neurospora crassa,* three checkpoints that regulate cell fusion at different stages have been characterized: pre-contact, contact-fusion and post-fusion systems (Goncalves *et al*. 2020). At the pre-contact checkpoint, chemotropic interactions between cells/hyphae are regulated by the “determinant of communication” or *doc* locus. This checkpoint ensures that cells/hyphae with *doc* alleles of identical specificity undergo chemotropic interactions that result in cell fusion. If cells have different allelic specificity at the *doc* locus, chemotropic interactions are disrupted, and cell fusion is rare (Heller *et al*. 2016; Goncalves *et al*. 2020). After chemotropic interactions and upon cell/hyphal contact, the next checkpoint is mediated by the “cell wall remodeling” or *cwr* locus, which is composed of two linked genes, *cwr-1* and *cwr-2*. If cells/hyphae have different allelic specificity at *cwr-1/cwr-2,* cell fusion is blocked at the cell wall dissolution stage (Supplementary Figure S1) (Goncalves *et al*. 2019; Detomasi *et al*. 2022). The third checkpoint functions post-fusion and results in regulated cell death of the fusion cells/hyphae (Rico-Ramirez *et al*. 2022). Genetic differences at post-fusion loci, such as *rcd-1* “regulator of cell death”, which is a homolog of mammalian gasdermin (Daskalov *et al*. 2019; Daskalov *et al*. 2020), *sec-9/plp-1* (Heller *et al*. 2018; Rico-Ramirez *et al*. 2022), or *het* loci (Saupe 2000; Glass and Kaneko 2003; Goncalves *et al*. 2017) triggers rapid vacuolization and cell death of the fusion cells or hyphal compartments. Previous calculations of allorecognition systems described in *N. crassa* that regulate cell fusion estimate ∼2,000,000 different genotypes in recombining populations, ensuring fusion only between near genetically identical individuals (Goncalves *et al*. 2020).

Phylogenetic analysis of sequences from a wild population of *N. crassa* (Ellison *et al*. 2011; Palma-Guerrero *et al*. 2013) revealed that the linked *cwr-1* and *cwr-2* alleles exhibit severe linkage disequilibrium and fall into six discrete haplogroups (HGs) (Supplementary Figure S2). Within each HG, the protein sequence of *cwr-1* and *cwr-2* alleles are completely conserved (Goncalves *et al*. 2019). Other filamentous ascomycete fungi, such as *Fusarium fujikuroi, F. verticillioides* and *N. discreta*, also have linked *cwr-1/cwr-2* loci that fell into discrete HGs in population samples (Goncalves *et al*. 2019). In *N. crassa*, allorecognition triggered by genetic differences at the *cwr* loci functions *in trans*, such that a cell/hyphae containing *cwr-1* from one HG shows a cell fusion block when in contact with a cell/hyphae containing *cwr-2* from a different HG (Supplementary Figure S1a) (Goncalves *et al*. 2019; Detomasi *et al*. 2022). Cells/hyphae that express *cwr-1* and *cwr-2* alleles from the same HG complete cell fusion. Strain bearing deletions of *cwr-1* and *cwr-2* (*Δcwr-1 Δcwr-2)* lose allorecognition capacity and undergo cell fusion with strains from any of the six *cwr* HGs. These data indicate that the *cwr* allorecognition functions to negatively regulate cell wall breakdown and remodeling associated with cell fusion.

*cwr-1* encodes a lytic polysaccharide monooxygenase (PMO) domain classified in the CAZY (Carbohydrate-Active enZYmes) as Auxiliary Activity Family 11 (AA11) (Hemsworth *et al*. 2014), with a glycine/serine linker and a potential chitin-binding domain (X278) at the C-terminus. The X278 domain is found in other predicted AA11 proteins and GH18 chitinases (Hemsworth *et al*. 2014; Goncalves *et al*. 2019). CWR-1 also contains an N-terminal histidine residue that is part of the histidine brace required for the binding a copper atom (Phillips *et al*. 2011; Detomasi *et al*. 2022). The PMO domain of CWR-1 exhibits C1-oxiding activity on chitin and has been shown to be necessary and sufficient to confer allorecognition. However, mutations that abolish the catalytic activity of the PMO domain do not affect allorecognition and the cell fusion block, indicating that CWR-1 has a moonlighting function as an allorecognition locus (Detomasi *et al*. 2022). Strains that only have *cwr-1* from one HG are blocked in cell fusion when interacting *in trans* with a second cell/hyphae that contains only *cwr-2* from a different HG (Goncalves *et al*. 2019). *cwr-2* is predicted to encode a plasma membrane protein with eight transmembrane domains, including two domains of unknown function, DUF3433 (Pfam PF11915) (Goncalves *et al*. 2019). The aim of this study was to clarify regions of CWR-1 and CWR-2 that confer allelic specificity in allorecognition leading to the cell fusion block. To achieve this, various *cwr-1* and *cwr-2* chimeras were designed based on structural analyses, which revealed regions important for regulating CWR allelic specificity.

## Materials and methods

### Strains and culture conditions

Strains are listed in Supplementary Table S1 and have been deposited at the Fungal Genetics Stock Center (https://www.plantpath.k-state.edu/research-services/fungal-genetics-stock-center/). Strains were grown in Vogel’s minimal medium (VMM) (Vogel 1956), with 1.5 % agar added for solid medium along with the necessary supplements. L-histidine (L-histidine hydrochloride monohydrate, 98%, Acros Organics) was added at a final concentration of 0.5 mg/mL to support the growth of histidine-auxotrophic strains. For crosses Westergaard’s synthetic cross-medium was employed (Westergaard and Mitchell 1947).

### Recombinant DNA techniques and plasmid constructions

Genes were amplified using genomic DNA of *N. crassa* strains, FGSC 2489, P4471, D111 (FGSC 8871), JW258, JW228 (Ellison *et al*. 2011; Palma-Guerrero *et al*. 2013) (Supplementary Table S1) as a template. PCR reactions were performed in MiniAmp Plus Thermal Cycler using Q5 High-Fidelity DNA polymerase (2000 U/mL) (NEB) according to the manufacturer’s instructions. The DNA fragments amplified were purified using Monarch DNA Gel extraction Kit (T1020S, NEB). Primers used in this study are reported in the Supplementary Table S2. All plasmids used are listed in Supplementary Table S3. All constructions were performed using Gibson Assembly^®^ Master Mix (E2611S, NEB) following the manufacturer’s instructions. The constructions generated were transformed into NEB 5-alpha Competent *E. coli* (NEB #C2987) for propagation and storage. The plasmid DNA was extracted with Monarch Plasmid Miniprep Kit (T1010S, NEB), in accordance with the manufacturer’s instructions.

The *cwr-1* HG1 or *cwr-1* HG6 chimeras were made using plasmid DNA *PMF::his-3::Pcwr-1-cwr-1-Tcwr-1* (from FGSC 2489; HG1) or *PMF::his-3-Ptef-1-cwr-1-Tcwr-1* (JW228; HG6) as a template, respectively. Chimera LC^HG1^ is under the regulation of the P*cwr-1^HG1^* promoter and chimera LC^HG6^ is under the regulation of the P*tef-1* promoter. The *cwr-2* HG1 chimeras were made using plasmid DNA *PMF::his-3-Ptef-1-cwr-2-V5-Tccg-1* (P4471; HG1) as a template for ED2 and ED4 domain swapping from *cwr-2* from a HG6 (JW228) strain. The *cwr-2* HG3 chimeras plasmid *PMF::his-3-Ptef-1-cwr-2-V5-Tccg-1* (JW258; HG3) as a template for swapping the ED2 and ED4 domains from *cwr-2* HG2 strain D111.

The *cwr-1^HG1^* and *cwr-1^HG1^* chimeric constructs were directed to the *his-3* locus (Margolin 1997) under the regulation of the native P*cwr-1^HG1^* promoter. The *cwr-1^HG6^*, *cwr-2^HG3^*and *cwr-2^HG6^* chimeric constructs and *cwr-1^HG1^, cwr-1^HG2^, cwr-1^HG3^, cwr-1^HG6^*, *cwr-2^HG1^*, *cwr-2^HG2^*, *cwr-2^HG3^* and *cwr-2^HG6^* constructs were targeted to *his-3* locus under the regulation of the P*tef-1* promoter. All chimeric strains and strains expressing a single *cwr-1* or *cwr-2* allele were constructed in the triple delete background *Δcwr-1; ΔNCU01381; Δcwr-2* (*ΔΔΔ*) (Goncalves *et al*. 2019); deletion of the NCU01381 gene has no influence on allorecognition (Goncalves *et al*. 2019). For the evaluation of the *cwr-2^HG1^* chimeras, strains expressing *cwr-1^HG1^* under the P*tef-1* promoter was used as one of the control strains. This strategy ensured that all strains expressing individual alleles and the *cwr-2^HG1^* chimeras were under the regulation of the same promoter. Supplementary Figure S3 compares cell fusion rates of strains expressing the *cwr-1^HG1^* allele under different promoters and at different loci, paired with either *cwr-2^HG1^* or *cwr-2^HG6^*; comparable fusion rates were obtained among all these strains.

### Transformation of Neurospora crassa, crosses and selection of homokaryons

*his-3* conidia from the *cwr* triple delete background were subjected to transformation with *Pac*I (R0547S, NEB), *Nde*I (R0111S, NEB) or *Ssp*I-HF (R3132S, NEB) linearized plasmids. Following established protocols (Margolin 1997), transformation was conducted through electroporation using a Bio-Rad Pulse controller plus and Bio-Rad gene pulser II. Electroporation was performed using 1 mm gap cuvettes (Bio-Rad Gene Pulser/MicroPulser Cuvette; Bio-Rad) at 1.5 kV, 600 ohm, 25 μF.

For each transformation, 30 histidine (His^+^) prototroph transformants were chosen and subsequently transferred to tubes containing VMM with Hygromycin B (10687010, 50 mg/mL; Thermo Fisher Scientific), at a final concentration of 200 μg/mL for selection. Integration of the constructs in the selected transformants was validated using the Phire Plant Direct PCR Kit (#F-130WH, Thermo Fisher Scientific) and Sanger sequencing. Selected heterokaryotic transformants were used as a female parent and cultivated on Westergaard’s synthetic crossing medium (Westergaard and Mitchell 1947) at 25 °C in constant light until the emergence of the protoperithecia. Strains were fertilized with a conidial suspension of the male parental strain, either FGSC 9716*/his-3* or FGSC 2489 *GFP/his-3 (his-3 csr-1::Pccg-1-gfp)* (Supplementary Table S1). Ascospores were subjected to heat shock at 60 °C for 40 minutes, followed by inoculation onto agar plates (VMM with 1.5% agar and mixture of 20% sorbose, 0.5% fructose, 0.5% glucose) and an overnight incubation at 30 °C. Germinated ascospores were transferred to VMM (Vogel 1956) slants. Homokaryotic selections were made based on histidine (His^+^) prototrophy, resistance to Hygromycin B (Hyg^+^), and/or to Cyclosporin A (Cyclosporin^+^). Cyclosporin A (30024, Sigma) was employed at a final concentration of 5 μg/mL for selection purposes. PCR analysis was performed to confirm the presence of the construct using the Phire Plant Direct PCR Kit (#F-130WH, Thermo Fisher Scientific) and subsequent Sanger sequencing.

### RNA isolation and cDNA synthesis

To extract RNA from the strains FGSC 2489, D111 (FGSC 8871), JW258, JW228 (Supplementary Table S1), 1×10^8^ conidia of each strain were resuspended in 100 mL of liquid VMM (Vogel 1956) and grown under constant light at 30 °C, 220 rpm, for 4 hr. Germlings were filtered using nitrocellulose membranes (0.45 µm) (#88018 Thermo Fisher Scientific), then added into 2 mL screw-cap tubes with 0.25g of 0.5 mm Zirconia/silica beads (BioSpec Products 11079105Z, #NC0450473, Fisher scientific) on dry ice. 1 mL TRIzol (#15596026, Thermo Fisher Scientific) was added to the frozen mycelia. Tubes were bead-beated at maximum speed for 1 min, 0.2 mL chloroform was added and vortexed. Subsequently, the samples were centrifugated for 15 min at 20000g and 4°C. 450 µL of the supernatant was transferred to a new Eppendorf tube, and 500 µL of isopropanol was added. The samples were gently shaken for 10 min at room temperature, followed by another centrifugation step at 20000g, 4°C for 10 min. The supernatant was discarded, and the pellet was washed with 75% ethanol, followed by another centrifugation step. The supernatant was discarded, and the pellet was dried and then resuspended in 87.5 µL of RNAse-free water and incubated at 60°C for 10 min. The samples were submitted to DNase digestion using RNase-Free DNase Set (#79254, QIAGEN), following the manufacturer’s instructions. Subsequently the RNA was cleaned using RNeasy Mini Kit (#74104, QIAGEN). For cDNA synthesis was used the ProtoScript^®^ II First Strand cDNA synthesis kit (E6560S, NEB).

### Germling-fusion assays and microscopy analysis

An aliquot of 200 μL of fresh conidia were stained with 40 μL of FM4-64 (styryl dye N-(3-triethylammoniumpropyl)–4-(6-(4-(diethylamino) phenyl) hexatrienyl) pyridinium dibromide, 514/670 nm absorption/emission, at a final concentration of 16.5 μM in ddH_2_O from a stock solution in DMSO of 16.5 mM (Invitrogen). Conidia were incubated a room temperature for 15 min in the dark and centrifuged in a microcentrifuge at 5000 rpm for 2 min, the supernatant was discarded, conidia were resuspended in 1 mL ddH_2_O and centrifuged; the procedure was repeated two times. The pellet was resuspended in 100 μL of sterile ddH_2_O and the number of conidia was counted using a hematocytometer. The concentration of the conidial suspension was adjusted to 3×10^7^ conidia/mL. An aliquot of 45 μL of the conidial suspension stained with FM4-64 was mixed with 45 μL of the conidia from strains expressing cytoplasmic GFP. 80 μL was plated on VMM agar plates (60 mm × 15 mm) and subsequently incubated for 3.5 hr at 30°C.

Agar rectangles measuring approximately 3 cm × 2 cm were excised and examined using a ZEISS Axioskop 2 MOT microscope. Images were taken employing a Q IMAGING FAST1394 COOLED MONO 12 BIT microscope camera (RETIGA 2000R SN:Q31594, 01-RET-2000R-F-M-12-C) with a Ph3 ×40/1.30 ∞/0.17 Plan-Neofluar oil immersion objective and processed using iVision-Mac Scientific Image Processing Bio Vision Technologies (iVision 4.5.6r4). The cell fusion percentage of germling pairings was determined by examining the cytoplasmic merging of GFP into germlings stained with FM4-64.

Images were captured across a minimum of 15 fields (for each DIC, Blue, Green, Red filter), documenting at least 100 germling-contact events, with three biological replicates. All images underwent further processing using Fiji (ImageJ2, version 2.14.0/1.54f; Build: c89e8500e4) (Schindelin *et al*. 2012). ANOVA statistical analyses were conducted using GraphPad Prism 10 (version 10.2.2) to identify differences in data across various strains and genotypes. A one-way ANOVA was applied when examining a single independent variable, whereas a two-way ANOVA was used for analyses involving two independent variables. Tukey’s test or Šídák’s test were subsequently performed for multiple comparisons to pinpoint significant differences. The p-values and 95% confidence intervals for each analysis are provided in the Supplementary Table S4. The images and figures were edited using Adobe Photoshop 2023 (Version 23.5.5).

### Prediction of protein structure and analysis using ColabFold

ColabFold V1.5.5 AlphaFold2 (Jumper *et al*. 2021; Mirdita *et al*. 2022), an online software that uses MMseqs2 (Many-against-Many sequence searching) to predict protein structures, was utilized to predict the three-dimensional structures of: PMO domain from the six different HGs, the CWR-1 chimeras and CWR-2 from HG1, HG2, HG3, HG6 and CWR-2 chimeras. (https://colab.research.google.com/github/sokrypton/ColabFold/blob/main/AlphaFold2.ipynb). For visualization of the predicted structures and their structural superposition, the PyMOL Molecular Graphics System Version 2.5.4 (Copyright (C) Schrödinger, LLC) was used. The Root Mean Square Deviation (RMSD) values were obtained using the align plugin in the PyMOL program (https://pymolwiki.org/index.php/Align), with default settings applied: 5 cycles of outlier rejection and a cutoff of 2.

### Illustrations

Digital illustrations were created using Adobe Illustrator 2023.

## Results

### The PMO LC domain is the most variable region between the six CWR-1 haplogroups

The PMO structure of the six different HGs was modeled using ColabFold (Mirdita *et al*. 2022), which showed a slightly different structure of the variable regions of the PMO domain among the six different HGs compared to previous SwissProt predictions (Detomasi *et al*. 2022). A comparison of the predicted three-dimensional PMO structures revealed two regions with the most structural variation among members of the different six CWR-1 HGs: L2 and LC (Figure 1a, b and c). Members within a CWR-1 HG showed no amino acid variation in either the L2 or LC regions (Goncalves *et al*. 2019) (Supplementary Figure S2). In the L2 region, there were noticeable conformational variations among the loops, particularly between amino acids 26 and 36 (PCQNTGGGY, using HG1 as a reference) (Figure 1a). In contrast, the LC region shows more pronounced structural differences in the loops, as well as variations in the positioning of the β-sheet secondary structure. Notably, only the PMO domain from HG1 and HG2 strains retain an α-helix at the end of the LC region (Figure 1a).

**Figure 1.**
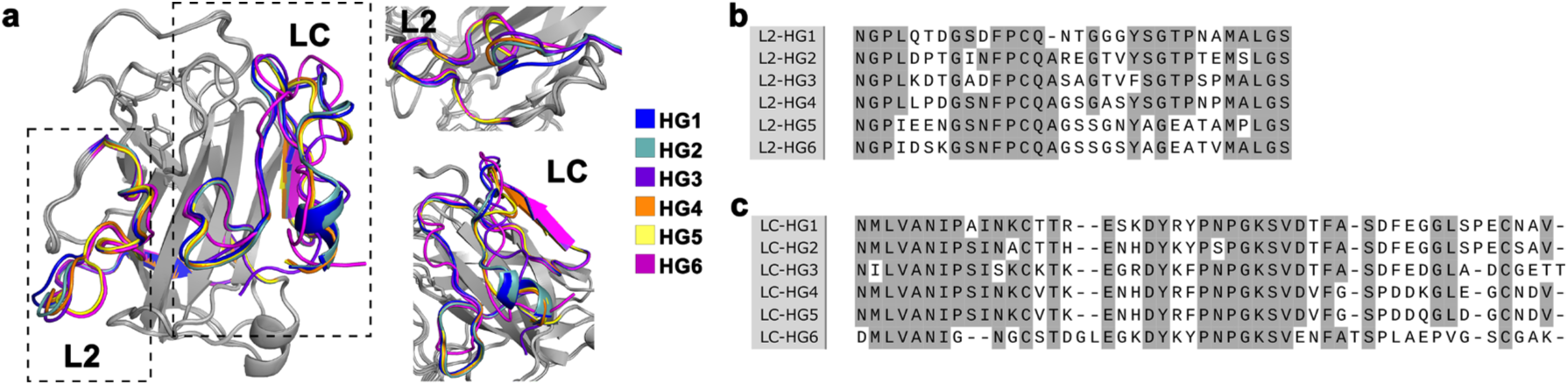
Model of the polysaccharide monooygenase (PMO) catalytic domain of CWR-1 from a member from each of the six different haplogroups (HGs). **a.** The models identify structural differences between members of the different CWR-1 HGs, specifically the L2 and LC regions. The two panels on the left display different views of these two regions. **b.** Alignment of representative sequences from one member of each of the six different HGs focused on the L2 region. c. Alignment of the amino acid sequences corresponding to LC region, using a representative sequence from a member from each HG. Conserved amino acids are highlighted in gray. The sequences used for modeling and alignment were from isolates FGSC 2489 (HG1), D111 (HG2), JW258 (HG3), JW242 (HG4), P4476 (HG5), and JW228 (HG6) (Ellison *et al*. 2011; Palma-Guerrero *et al*. 2013). These sequences are representative of members of each of the respective HGs. No significant amino acid sequence differences are present in CWR-1 between members of the same HG (Goncalves *et al*. 2019). Alignment was performed using MAFFT version 7 (https://mafft.cbrc.jp/alignment/server/) and edited with SnapGene Viewer (Version 7.2.1). Prediction of crystal structure analysis using ColabFold V1.5.5 AlphaFold2 (Jumper *et al*. 2021; Mirdita *et al*. 2022).

To assess the similarities between the different conformational structures of the PMO domains from the six different CWR-1 HGs, quantitative data were obtained by calculating the Root Mean Square Deviation (RMSD). The RMSD provides a numerical measure of the structural similarity between two aligned atomic configurations, expressed in angstroms (Å) (Kabsch 1976; Kufareva and Abagyan 2012). Values closer to zero indicate higher similarity, while larger values suggest greater structural deviations. Supplementary Table S5 shows the results of the comparisons of the conformational structures of the PMO domain from each of the six HGs. With overall RMSD values ranging from 0.171 Å to 0.281 Å, the deviations are minimal, suggesting only slight structural differences. For specific regions, the L2 and LC regions show slightly higher variability (0.208 Å to 0.448 Å and 0.151 Å to 0.494 Å, respectively), but still within a range that signifies strong similarity. RMSD values below 1 Å are generally considered very similar (Chothia and Lesk 1986), indicating minimal conformational differences across the PMO domain HG structures.

We first assessed fusion percentages between strains with wild type *cwr-1* and *cwr-2* alleles from the most divergent HGs (HG1 and HG6; Supplementary Figure S2). These *cwr-1/cwr-2* alleles from members of the different HGs were targeted to the *his-3* locus in the triple-deletion strain, *Δcwr-1 ΔNCU01381 Δcwr-2* (*ΔΔΔ*). This mutant (*ΔΔΔ*) is capable of undergoing cell fusion with both *cwr* compatible and incompatible strains due to the absence of the genes responsible for triggering allorecognition at the cell fusion checkpoint (Goncalves *et al*. 2019). Fusion assays were conducted using one strain expressing cytoplasmic GFP and the partner strain stained with the FM4-64 (Hickey *et al*. 2002; Hickey *et al*. 2004). If cell fusion occurred, GFP migrated to the cell stained with the FM4-64, resulting in a pink tint against a green background. However, if no cell fusion occurred, germlings were observed in physical contact, but transfer of GFP fluorescence did not occur (Figure 2a).

**Figure 2.**
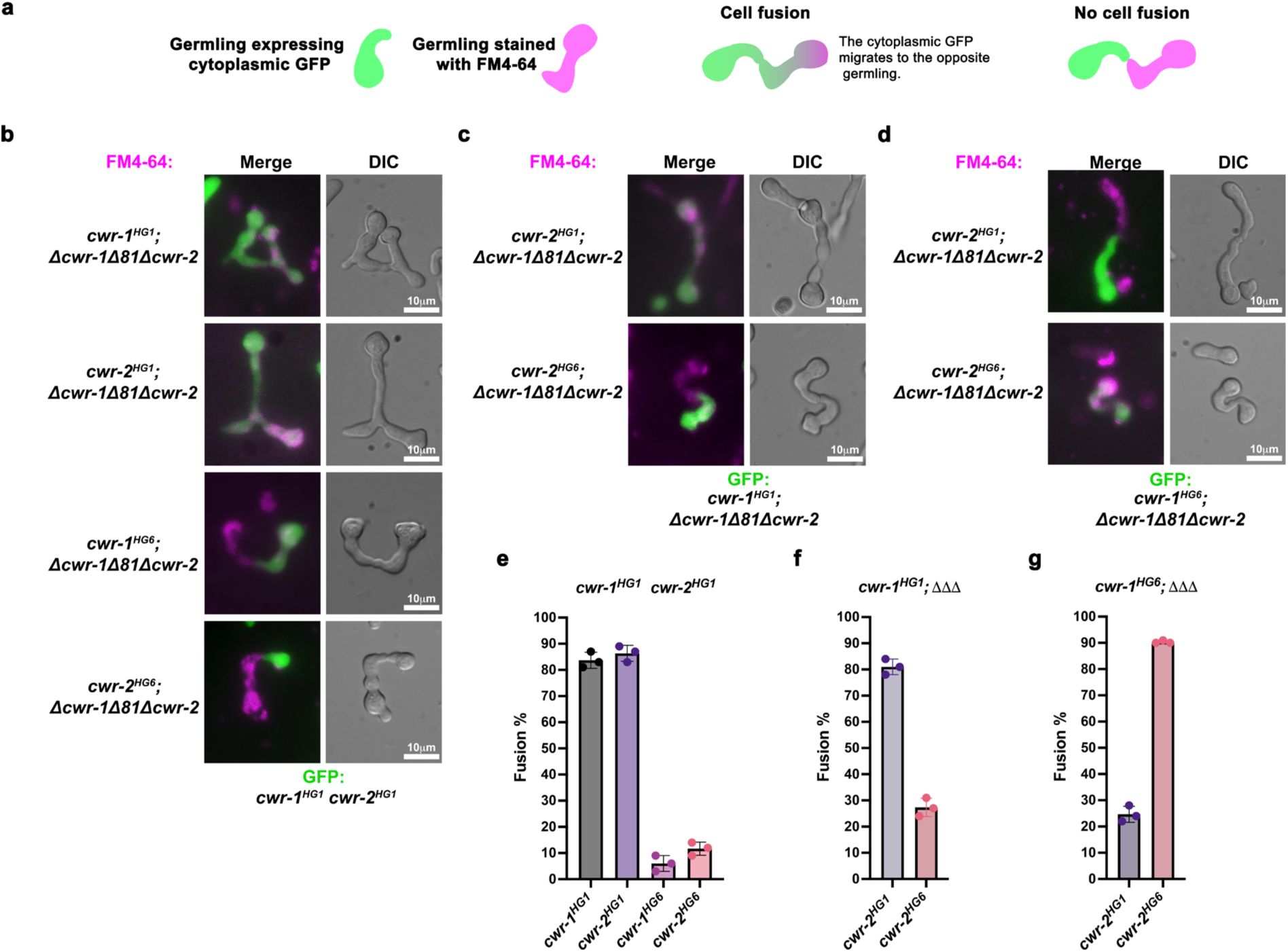
*cwr-1* and *cwr-2* from different HGs function in *trans* to trigger allorecognition and fusion block at the cell wall remodeling checkpoint. **a.** Diagram to show experimental design to test cell fusion percentages where germlings from one HG express cytoplasmic GFP while germlings from another HG are stained with the dye FM4-64 (left panel). If germlings contain compatible *cwr-1* and *cwr-2* alleles, germlings undergo cell fusion, and the migration of cytoplasmic GFP protein into the opposite germling stained with FM4-64 is observed (middle panel). If germlings contain incompatible *cwr-1* and *cwr-2* alleles, allorecognition is triggered and cell fusion is blocked; no transfer of GFP between germlings is observed (right panel). **b**. Micrographs showing the most common observed events in fusion tests. Germlings from a wild type HG1 strain (*cwr-1^HG1^* and *cwr-2^HG1^)* expressing cytoplasmic GFP paired with germlings harboring *cwr-1^HG1^* or *cwr-2^HG1^*(compatible combinations) or with germlings harboring *cwr-1^HG6^*or *cwr-2*^HG6^ (incompatible combinations). All strains were constructed in the triple delete background *Δcwr-1; ΔNCU01380; Δcwr-2* (*ΔΔΔ*) and the *cwr-1* or *cwr-2* alleles were targeted to the *his-3* locus. **c**. Micrographs showing fusion tests between a strain bearing *cwr-1^HG1^; ΔΔΔ* paired with germlings that were either *cwr-2^HG1^; ΔΔΔ* or *cwr-2^HG6^; ΔΔΔ.* d. Micrographs showing fusion tests between germlings carrying *cwr-1^HG6^; ΔΔΔ* paired with *cwr-2^HG1^; ΔΔΔ* or *cwr-2^HG6^; ΔΔΔ* germlings. e. Quantification of cell fusion events shown in panel B. **f**. Quantification of cell fusion events shown in panel C. **g**. Quantification of cell fusion events shown in panel D. Fusion tests were performed in biological triplicate, assessing fusion interactions of 100 germling pairs for each replicate. Individual p-values are reported in Supplementary Table S4.

To evaluate cell fusion percentages of strains bearing different *cwr-1* or *cwr-2* alleles, we first paired them with a HG1 strain (FGSC 2489), which carries both *cwr-1^HG1^* and *cwr-2^HG2^* alleles at the native locus. When the *cwr-1^HG1^ cwr-2^HG1^* strain was paired with strains bearing either *cwr-1^HG1^* or *cwr-2^HG1^*, high cell fusion percentages were observed, reaching 83% and 86%, respectively (Figure 2e). These data indicated that the engineered strains were fusion proficient. We then assessed fusion in strains bearing the most distantly related *cwr-1/cwr-2* alleles: HG1 paired with HG6 (Supplementary Figure S2). In contrast to fusion percentages between strains bearing *cwr-1* and *cwr-2* from the same HG, fusion percentages of *cwr-1^HG1^ cwr-2^HG1^ + cwr-1^HG6^* and *cwr-1^HG1^ cwr-2^HG1^* + *cwr-2^HG6^*pairings were very low (6% and 11%, respectively; Figure 2b and e).

In wild isolates, allorecognition is triggered by two non-allelic *cwr* interactions: *cwr-1* in cell 1 with *cwr-2* in cell 2 and *cwr-2* in cell 1 with *cwr-1* in cell 2 (Supplementary Figure S1a), potentially leading to a more robust allorecognition response. We therefore engineered strains with *cwr-1* or *cwr-2* alleles targeted to the *his-3* locus in the *ΔΔΔ* background; these engineered strains were completely isogenic except for *cwr-1* or *cwr-2.* In pairings between strains with compatible alleles, such as a *cwr-1^HG1^* + *cwr-2^HG1^* or *cwr-1^HG6^* + *cwr-2^HG6^* pairings, a high percentage of fusion was observed (81% and 90%, respectively) (Figure 2c, d, f and g). However, pairings between *cwr-1^HG1^* + *cwr-2^HG6^* strains showed a fusion percentage of 27% (Figure 2c and f; Supplementary Figure S3b), which was a significantly higher fusion percentage than of *cwr-1^HG1^ cwr-2^HG1^* + *cwr-2^HG6^* pairings (11%). Similarly, *cwr-1^HG6^* + *cwr-2^HG1^* pairings showed a fusion percentage of 24%, also a significantly higher fusion percentage than pairings between *cwr-1^HG1^ cwr-2^HG1^* + *cwr-1^HG6^* strains (6%) (Figure 2d, e and g; Supplementary Figure S1c).

To determine whether fusion behavior was restricted to pairings between HG6 and HG1 strains, we also assessed interactions between strains bearing *cwr* alleles from a HG2 and HG3 strains; *cwr-1/cwr-2* in HG2 and HG3 are more closely related than HG1 and HG6 alleles (Supplementary Figure S2). Pairings between *cwr-1^HG3^* + *cwr-2^HG2^* strains showed a fusion percentage of 52%, while *cwr-1^HG2^* + *cwr-2^HG3^* pairings exhibited a fusion percentage of 24% (Supplementary Figure S1b). These observations indicate that one *cwr-1/cwr-2* HG non-allelic combination may confer a greater degree of fusion block than other combinations (Supplementary Figure S1b, c). In pairings with the *cwr-1^HG1^ cwr-*2^HG1^ (FGSC 2489) strain, fusion percentages were reduced, with a fusion rate of 14% between *cwr-1^HG1^ cwr-*2^HG1^ + *cwr-1^HG2^*, 8% in pairings between *cwr-1^HG1^ cwr-2^HG1^* + *cwr-2^HG2^*, 15% in pairings between *cwr-1^HG1^ cwr-*2^HG1^ + *cwr-1^HG3^* and 8% in pairings between *cwr-1^HG1^ cwr-*2*^HG1^* + *cwr-2^HG3^* (Supplementary Figure S1d). These data indicated that strains that contained only one *cwr* non-allelic interaction did not show as robust allorecognition response as in pairings with a wild type strain, where *cwr-1* and *cwr-2* are at their native locus. It is possible that *cwr-1* and *cwr-2* at the native locus functions better than targeted insertions of *cwr-1* or *cwr-2* at the *his-3* locus, or that interactions between CWR-1 and CWR-2 within the same cell may be playing a role to enhance the cell fusion block.

### Examining the impact of the L2 and LC regions on allorecognition at the cell fusion checkpoint

We hypothesized that the L2 and LC regions that were divergent between *cwr-1* haplogroups, but identical within members of the same haplogroup, would be involved in *cwr* allelic specificity (Figure 1b, c). To test this hypothesis, chimeras were constructed using *cwr-1* from HG1, where the L2 (H35-A61) and LC (N182-V229) regions of the PMO domain were exchanged with their homologous counterparts from HG6 (L2: Q35-S67, LC: D183-K230). Three chimeras *cwr-1^HG1^L2^HG6^, cwr-1^HG1^LC^HG6^*and *cwr-1^HG1^L2^HG6^LC^HG6^* were constructed (Figure 3a). The structural conformation of the chimeric PMO domains were also assessed using ColabFold (Mirdita *et al*. 2022); the three-dimensional structures displayed a conformation very similar to the predicted PMO^HG1^ domain but with notable variations in the L2^HG6^ and LC^HG6^ regions (Figure 3b). Strains carrying the three different chimeras, *cwr-1^HG1^L2^HG6^, cwr-1^HG1^LC^HG6^* or *cwr-1^HG1^L2^HG6^LC^HG6^*, were first tested for cell fusion with the triple deletion strain (*ΔΔΔ*); all three chimeras showed robust cell fusion percentages (∼82%), validating that the constructs did not disrupt the cell fusion process (Figure 3c).

**Figure 3.**
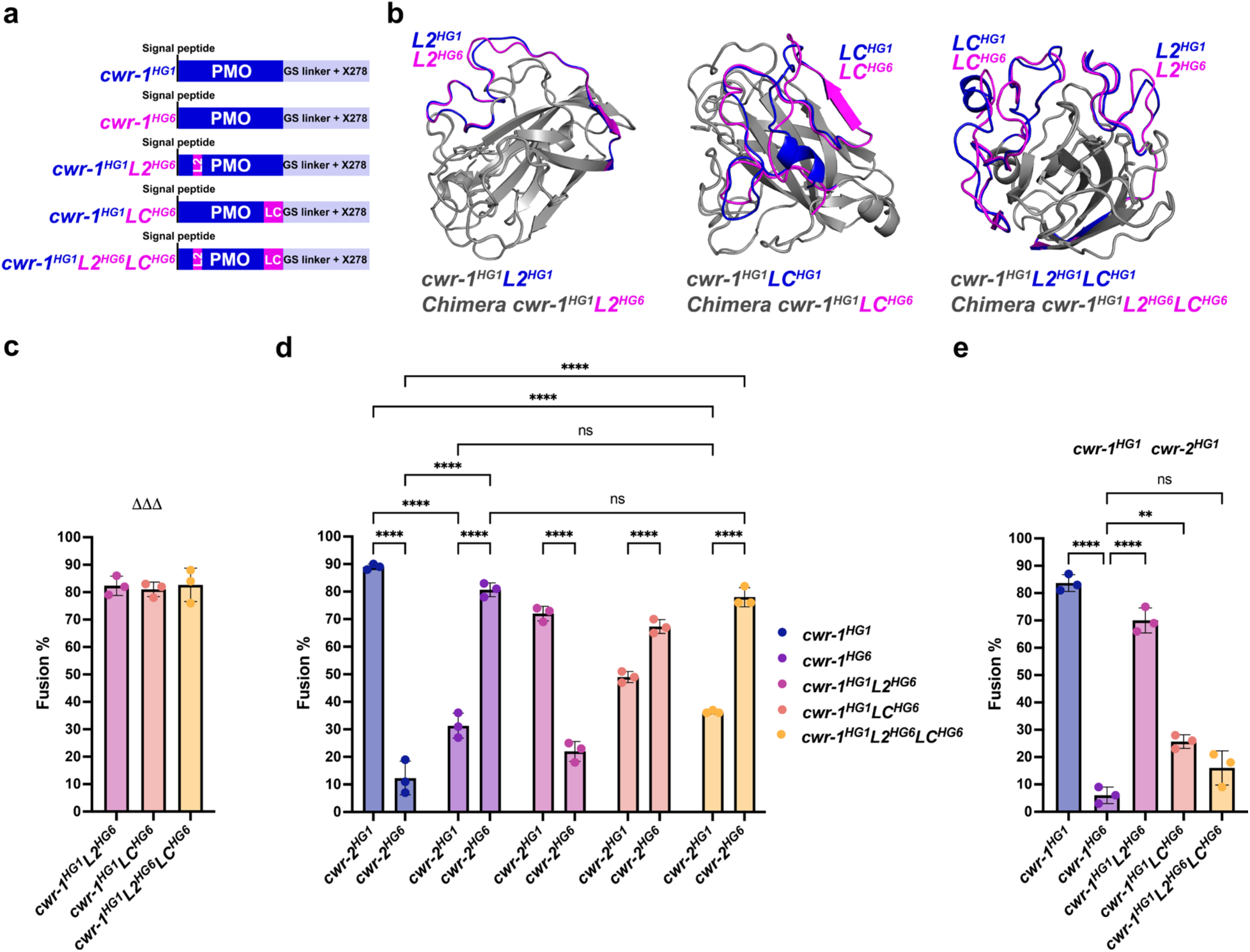
Evaluation of the effects on allorecognition at the cell fusion check point of the CWR-1^HG1^ PMO chimeras, with L2^HG6^ or LC^HG6^ or L2^HG6^/LC^HG6^. **a**. Schematic of CWR-1 HG1/HG6 chimeric constructs. The PMO (polysaccharide monooxygenase) domain, GS linker (glycine, serine linker) and X278 domain (predicted chitin binding domain) are indicated. The L2 or LC regions, or both, of the CWR-1^HG1^ PMO domain were replaced by the corresponding regions from the PMO domain of HG6. **b**. The three panels show the predicted protein structures of the HG1/HG6 CWR-1 chimeras (*cwr-1^HG1^L2^HG6^, cwr-1^HG1^LC^HG6^* and *cwr-1^HG1^L2^HG6^LC^HG6^*) generated by ColabFold (Mirdita *et al*. 2022) and overlapped with the PMO of the CWR-1^HG1^ (grey). **c**. Quantification of cell fusion events between strains harboring the CWR-1^HG1^ chimeras (bearing L2, LC or L2/LC from a *cwr-1* HG6 allele) paired with a strain lacking both *cwr-1* and *cwr-2* (*ΔΔΔ*). All chimeric constructs were targeted to the *his-3* locus in the *ΔΔΔ* background. **d**. Quantification of cell fusion events between strains harboring the CWR-1^HG1^ chimeras (bearing L2, LC or L2/LC from a *cwr-1* HG6 allele) paired with germlings harboring either *cwr-2^HG1^* or *cwr-2^HG6^*alleles in the *ΔΔΔ* background. A two-way ANOVA followed by Tukey’s post-hoc test was used for statistical analysis, error bars represent SD (Standard deviation), ****p<0.0001, ns: not significant. **e**. Quantification of cell fusion events between strains harboring the CWR-1^HG1^ chimeras (bearing L2, LC or L2/LC from a *cwr-1* HG6 allele) paired with *cwr-1^HG1^ cwr-2^HG1^* germlings (FGSC 2489). As controls, germlings expressing either *cwr-1^HG1^* or *cwr-1^HG6^* were paired with *cwr-1^HG1^ cwr-2^HG1^* germlings (FGSC 2489). A one-way ANOVA followed by Tukey’s post-hoc test was used for statistical analysis, error bars represent SD (Standard deviation), **p<0.01, ****p<0.0001, ns: not significant. The cell fusion test experiments were performed in biological triplicate, assessing fusion of 100 germling pairs for each replicate. Individual p-values are reported in Supplementary Table S4.

Strains expressing the *cwr-1^HG1^L2^HG6^* + *cwr-2^HG1^* pairings showed a slight reduction in cell fusion percentages (72%), as compared to cell fusion percentages of 89% in control pairings (*cwr-1^HG1^* + *cwr-2^HG1^*) (Figure 2d). The *cwr-1^HG1^LC^HG6^* + *cwr-2^HG1^* pairings displayed a more pronounced reduction in cell fusion (51%). Swapping both L2 and LC (*cwr-1^HG1^L2^HG6^LC^HG6^* + *cwr-*2^HG1^) pairings resulted in a more significant reduction in cell fusion (36% fusion) and which was indistinguishable from the control pairings between *cwr-1^HG1^*+ *cwr-2^HG6^* strains (Figure 3d). Significantly, both the *cwr-1^HG1^LC^HG6^* + *cwr-2^HG6^*and *cwr-1^HG1^L2^HG6^LC^HG6^* + *cwr-2^HG6^*pairings showed a significant increase in fusion percentages (67% and 78%, respectively) (Figure 3d). Fusion percentages of the *cwr-1^HG1^L2^HG6^LC^HG6^* + *cwr-2^HG1^* and *cwr-1^HG1^L2^HG6^LC^HG6^* + *cwr-2^HG6^* pairings were indistinguishable from the control pairings between *cwr-1^HG6^* + *cwr-2^HG1^* and *cwr-1^HG6^* + *cwr-2^HG6^*. Thus, swapping the L2 and LC region from a HG6 strain into the PMO domain of a HG1 strain switched its specificity to a HG6 strain. These data strongly support the involvement of the L2 and particularly the LC region in *cwr-1* allelic specificity and allorecognition.

The fusion percentages of the *cwr-1^HG1^L2^HG6^, cwr-1^HG1^LC^HG6^* and *cwr-1^HG1^L2^HG6^LC^HG6^*chimeric strains were also assessed when paired with the *cwr-1^HG1^ cwr-2^HG1^* strain (FGSC 2489). The *cwr-1^HG1^ cwr-2^HG1^* + *cwr-1^HG1^L2^HG6^* chimera showed a slightly reduced fusion percentage (70%) as compared to the fusion percentage between *cwr-1^HG1^ cwr-2^HG1^*+ *cwr-1^HG1^* strains (84%) (Figure 2e). However, pairings between the *cwr-1^HG1^ cwr-2^HG1^* + *cwr-1^HG1^LC^HG6^*showed a significant reduction in fusion percentages (26%), while the *cwr-1^HG1^ cwr-2^HG1^* + *cwr-1^HG1^L2^HG6^LC^HG6^* pairings displayed and even more reduced fusion (16%).

To determine whether the *cwr-1* L2 and LC allelic swaps were symmetrical in function, chimeras were constructed with a *cwr-1^HG6^* allele with the L2^HG6^ (Q35-S67) and LC^HG6^ (D183-K230) regions replaced with L2^HG1^ (H35-S66) and LC^HG1^ (N182-V229) regions from the PMO domain of *cwr-1^HG1^*(Figure 4a). The chimeric proteins were modeled, and the overlapping analysis showed that swapping the two motifs, L2^HG1^ and LC^HG1^, did not significantly affect the conformational structure of *cwr-1^HG6^* PMO domain (Figure 4b). The three chimeras (*cwr-1^HG6^L2^HG1^*; *cwr-1^HG6^LC^HG1^*; *cwr-1^HG6^L2^HG1^ LC^HG1^*) were first tested for cell fusion by pairing them with the triple-deletion strain and all showed high percentages of cell fusion (Figure 4c).

**Figure 4.**
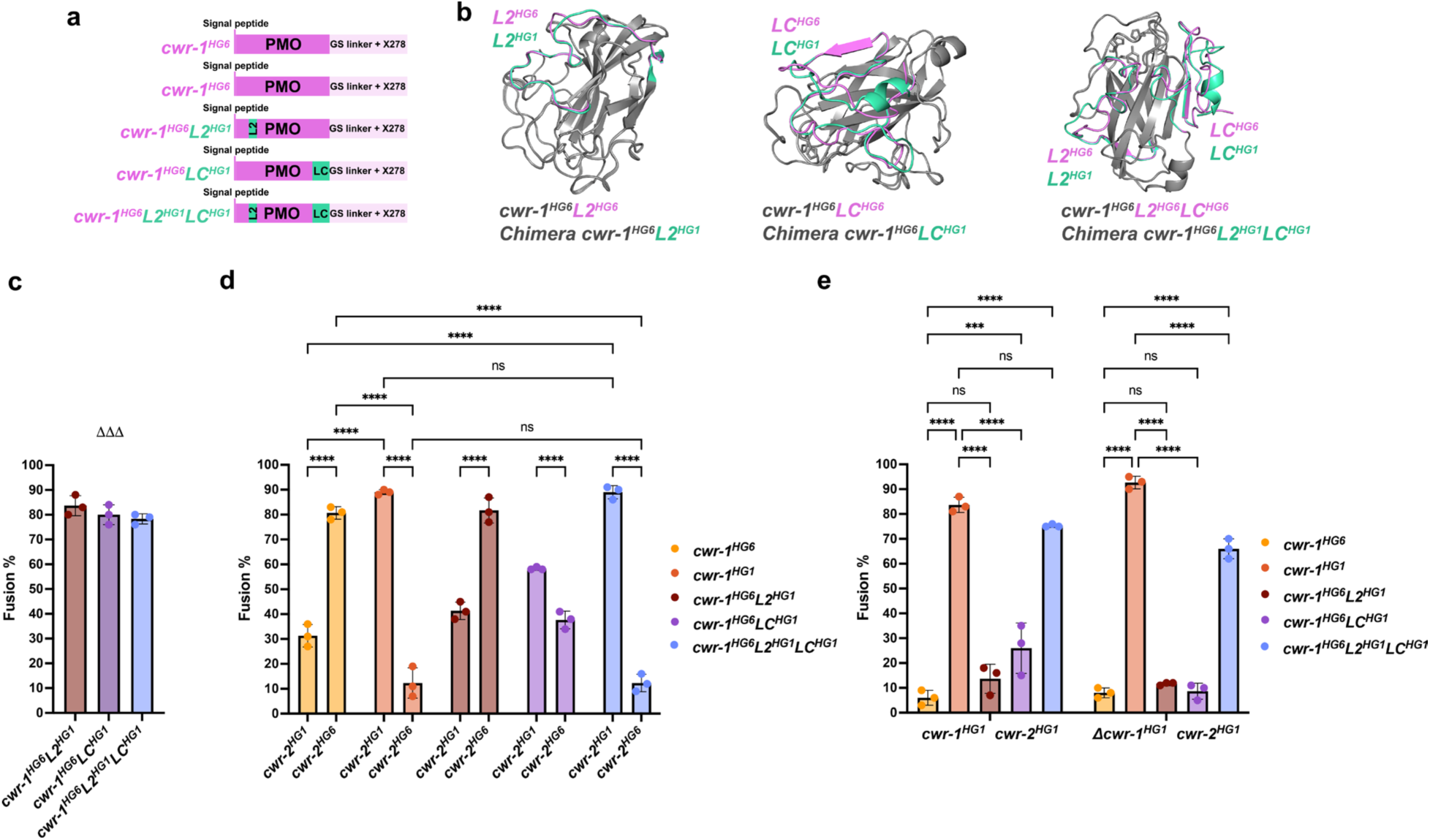
Evaluation of the effects on allorecognition and cell fusion at the cell wall remodeling checkpoint of the CWR-1^HG6^ PMO chimeras. **a**. Schematic of the CWR-1^HG6^ L2^HG1^, LC^HG1^ and L2^HG1^/LC^HG1^ chimeras. PMO (polysaccharide monooxygenase domain), GS linker (glycine, serine linker domain) and X278 domain (predicted chitin binding domain) are indicted. **b**. The three panels show the predicted structures of the CWR-1^HG6^L2^HG1^, CWR-1^HG6^LC^HG1^ and CWR-1^HG6^L2^HG1^LC^HG1^ chimeric proteins generated by ColabFold (Mirdita *et al*. 2022) and overlapped with the PMO of the CWR-1^HG6^ (grey). **c**. Quantification of cell fusion events of *cwr-1*^HG6^L2^HG1^*, cwr-1*^HG6^LC^HG1^ or *cwr-1*^HG6^L2^HG1^LC^HG1^ strains paired with a strain lacking both *cwr-1* and *cwr-2* (*ΔΔΔ*). All chimeric constructs were targeted to the *his-3* locus in the *ΔΔΔ* background. **d**. Quantification of cell fusion events between germlings carrying the *cwr-1*^HG6^L2^HG1^*, cwr-1*^HG6^LC^HG1^ or *cwr-1*^HG6^L2^HG1^LC^HG1^ chimeras paired with germlings bearing *cwr-2^HG1^* or *cwr-2^HG6^*alleles targeted to the *his-3* locus in the *ΔΔΔ* background. Two-way ANOVA followed by Tukey’s post-hoc test was used for statistical analysis, error bars represent SD (Standard deviation), ****p<0.0001, ns: not significant. **e.** Quantification of fusion events of germlings carrying the *cwr-1*^HG6^L2^HG1^*, cwr-1*^HG6^LC^HG1^ or *cwr-1*^HG6^L2^HG1^LC^HG1^ chimeras with the *cwr-1^HG1^ cwr-2^HG1^* strain (FGSC 2489) or *Δcwr-1^HG1^ cwr-2^HG1^* germlings. A Two-way ANOVA followed by Tukey’s post-hoc test was used for statistical analysis, error bars represent SD (Standard deviation), ***p<0.001, ****p<0.0001, ns: not significant. Cell fusion tests were performed in biological triplicate, assessing fusion of 100 germling pairs for each replicate. Individual p-values are reported in Supplementary Table S4.

The *cwr-1^HG6^L2^HG1^* + *cwr-2^HG1^* pairings showed a fusion percentage of 41%, but a high fusion percentage of 82% in *cwr-1^HG6^L2^HG1^* + *cwr-2^HG6^*pairings (Figure 4d). In contrast, the *cwr-1^HG6^LC^HG1^* + *cwr-2^HG6^* pairings showed a fusion percentage of 38%, while *cwr-1^HG6^LC^HG1^* + *cwr-2^HG1^* pairings showed a fusion percentage of 58% (Figure 4d). Pairings between *cwr-1^HG6^L2^HG1^LC^HG1^* + *cwr-2^HG6^* showed a fusion percentage of only 12%, a value that was indistinguishable to that of control pairings between *cwr-1^HG1^* + *cwr-2^HG6^*. In *cwr-1^HG6^L2^HG1^LC^HG1^* + *cwr-2^HG1^* pairings, the fusion percentage was 89%, a value indistinguishable to the control pairings (*cwr-1^HG1^* + *cwr-2^HG1^*) (Figure 4d). These data indicated that swapping both the L2 and LC regions of *cwr-1^HG6^* with L2 and LC region from a *cwr-1^HG1^* strain completely switched *cwr-1* allelic specificity to a HG1 strain.

As additional controls, the *cwr-1^HG6^L2^HG1^*, *cwr-1^HG6^LC^HG1^* and *cwr-1^HG6^L2^HG1^LC^HG1^* chimeric strains were also paired with the wild type *cwr-1^HG1^ cwr-2 ^HG1^* strain and a mutant strain *Δcwr-1^HG1^ cwr-2^HG1^* that expresses *cwr-2^HG1^* from the native locus. As shown in Figure 4e, the *cwr-1^HG1^ cwr-2 ^HG1^* + *cwr-1^HG6^L2^HG1^LC^HG1^* and *Δcwr-1^HG1^ cwr-2 ^HG1^* + *cwr-1^HG6^L2^HG1^LC^HG1^* pairings showed high fusion percentages (75% and 66%), indicating that swapping the L2 and LC region of the PMO domain has an important role in allelic specificity of *cwr-1*. In the fusion tests with the engineered strains, the LC region has a more significant influence than the L2 region for switching allelic specificity (Figure 3d and Figure 4d). However, in fusion tests with the wild type (*cwr-1^HG1^ cwr-2 ^HG1^*) and the *cwr-1* deletion strain (*Δcwr-1^HG1^ cwr-2^HG1^)*, where the *cwr-2* alleles are at the native locus, both the L2 and LC regions were required for switching allelic specificity (Figure 4e).

### The LC region alone causes an alteration in allelic specificity at the cell fusion checkpoint

To assess whether the LC region alone can affect allorecognition at the cell wall remodeling checkpoint, the LC regions of HG1 and HG6 were first modeled independently using ColabFold, comparing each three-dimensional model with the corresponding predicted structure of the PMO domain for HG1 and HG6. The predicted structures of the LC regions for both HG1 and HG6 exhibited variations in their three-dimensional conformations compared to the LC structure within an intact PMO domain. These changes included the absence of a beta sheet (Figure 5a and b, arrows in green) and an alpha helix in the architecture of LC from the HG1 PMO domain (Figure 5a, pink arrow). Two chimeric constructs were designed, LC^HG1^ and LC^HG6^, by incorporating the signal peptide of CWR-1, the LC region of the PMO domain, the glycine and serine linker domain and the X278 domain (Figure 5c). The RMSD values were calculated for both chimeras (Kufareva and Abagyan 2012). The comparison of the three-dimensional structure of the LC HG1 chimera with PMO HG1 resulted in an RMSD of 1.306 Å, while the comparison of LC HG6 with PMO HG6 resulted in a RMSD of 1.305 Å (Supplementary Table S5). These values suggest that the chimeras retain a similar overall structure to the original PMO region, although with some differences in structural conformation. RMSD values around 1 Å generally indicate high structural similarity, with minor but noticeable conformational changes (Chothia and Lesk 1986).

**Figure 5.**
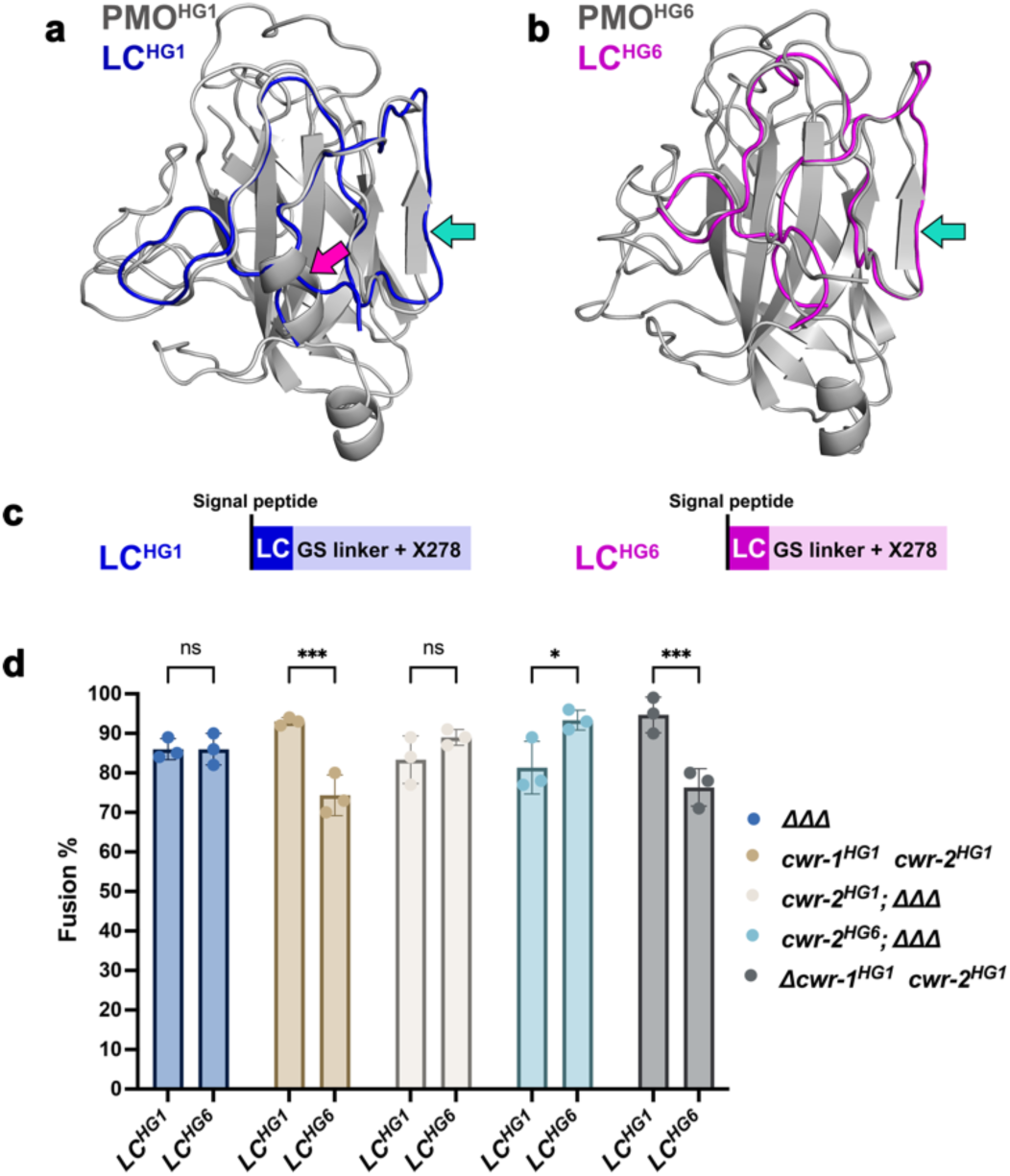
LC domain alone is not sufficient to trigger full allorecognition to block cell fusion. **a**. Overlap of the structural conformation between the PMO domain of HG1 and the LC domain HG1. **b**. Overlap of the structural conformation between the PMO domain of HG6 and the LC domain HG6. The predicted structures for a and b were generated by ColabFold (Mirdita *et al*. 2022). The pink arrow indicates the missing alpha helix in LC^HG1^, while the green arrow indicates the missing beta sheet in both LC structures. **c**. Schematic showing the LC^HG1^ and LC^HG6^ constructs. The LC^HG1^ and LC^HG6^ constructs have a deletion of the PMO domain and contain the GS linker and the predicted X278 chitin binding domain. All chimeric constructs are targeted to *his-3* locus. **d**. Quantification of fusion events of LC^HG1^ and LC^HG6^ chimeric strains paired with the strains indicated in the graphic. A two-way ANOVA followed by Šídák’s multiple comparison test was used for statistical analysis, error bars represent SD (Standard deviation), *p<0.05, ***p<0.001, ns: not significant. Cell fusion tests were performed in biological triplicate, assessing fusion events between 100 germling pairs for each replicate. Individual p-values are reported in Supplementary Table S4.

The LC^HG1^-GS-X278 and LC^HG6^-GS-X278 chimeric strains were paired with the *ΔΔΔ* mutant and fusion percentages were high (86%) (Figure 5d). Pairings between *cwr-1^HG1^ cwr-2^HG1^* + LC^HG1^-GS-X278 showed a 93% fusion percentage, while *cwr-1^HG1^ cwr-2^HG1^* + LC^HG6^-GS-X278 pairings showed a lower fusion percentage (74%). Pairings between *Δcwr-1^HG1^ cwr-2^HG1^* + LC^HG1^-GS-X278 pairings showed a fusion percentage of 95%, while *Δcwr-1^HG1^ cwr-2^HG1^* + LC^HG6^GS-X278 pairings had a fusion percentage of 76% (Figure 5d), a statistically significant difference. In LC^HG1^-GS-X278 + *cwr-2^HG6^* pairings, a significant difference (p<0.05) in fusion percentage was observed (81%) as compared to a 93% fusion percentage in LC^HG6^-GS-X278 + *cwr-2^HG6^* pairings. These results suggest that the LC region alone can trigger allorecognition and a cell fusion block to a degree, thus reducing cell fusion when paired with strains expressing incompatible *cwr-1* alleles.

### Assessing the effect of the ED2 and ED4 extracellular domains of CWR-2 on allorecognition at the cell fusion checkpoint

CWR-2 is a predicted transmembrane protein composed of four extracellular domains (ED), five cytoplasmic domains (CD), and eight transmembrane domains. It contains two domains of unknown function (DUF3433) (Supplementary Figure S4). We initially experienced difficulty in cloning *cwr-2* alleles from some of the haplogroups, suggesting that the annotation for the *cwr-2* ORF might be different in members of the different *cwr-2* HGs. We therefore isolated RNA from strains from different *cwr-2* HGs, which was subjected to RT-PCR to identify the *cwr-2* introns (see Materials and Methods). Comparative analysis of *cwr-2* sequences showed that members of HG1, HG2, and HG3 each contained three introns, while members in HG6 exhibited an additional intron positioned within the coding region for the extracellular domain 4 (ED4) (Supplementary Figure S5). CWR-2 is a large protein that differs in size among the CWR-2 haplogroups: 1217 amino acids (aa) in CWR-2^HG1^ isolates, 1220 aa in CWR-2^HG2^ isolates, 1216 aa in CWR-2^HG3^ isolates, 1218 aa in CWR-2^HG4^ isolates, 1233 aa in CWR-2^HG5^ isolates, and 1273 aa in CWR-2^HG6^ isolates. The spatial structure of CWR-2, as predicted by ColabFold (Mirdita *et al*. 2022), provided a clear depiction of the conformational structure in space of CWR-2 and its complexity, and shows that two of the extracellular domains, ED2 and ED4, are extensive and prominent in comparison to the ED1 and ED3 extracellular domains (Supplementary Figure S6). The transmembrane domains of CWR-2 are composed of alpha helices composed of approximately 21 amino acids. The cytoplasmic domains are composed of a combination of alpha helices, beta sheets, and loops and are arranged in close proximity, making it difficult to differentiate them. However, the CD5 domain in HG6 is notably larger compared to members of the other HGs, containing approximately 40 additional amino acids (Supplementary Figure S4).

To identify regions important for CWR-2 allelic specificity, we first examined members of the most distant haplogroups, *cwr-2*^HG1^ and *cwr-2*^HG6^ (Supplementary Figure S2). We hypothesized that ED2 and ED4 extracellular domains of CWR-2, due to their large size, sequence diversity and orientation towards the cell wall, might play a crucial role in allorecognition. The overlay of CWR-2^HG1^ and CWR-2^HG6^ structures revealed significant differences in these domains (Figure 6a). A structural comparison between the two CWR-2 proteins generated an RMSD value of 1.690 Å, indicating that CWR-2^HG1^ and CWR-2^HG6^ share similar three-dimensional structures, with some variations across the entire proteins. Analyzing the ED2 and ED4 domains separately resulted in RMSD values of 0.730 Å and 0.650 Å, respectively, indicating closer similarity in these regions compared to the entire CWR-2^HG1^ and CWR-2^HG6^ proteins. For the design of the *cwr-2^HG1/HG6^* chimeras, the extracellular domains ED2^HG1^ (G164-R496) and ED4^HG1^ (N746-A1100) were exchanged for the corresponding ED2^HG6^ (G164-R500) and ED4^HG6^ (F760-A1113) domains of CWR-2^HG6^, generating three different chimeras: *cwr-2^HG1^ED2^HG6^, cwr-2^HG1^ED4^HG6^,* and *cwr-2^HG1^ED2^HG6^ ED4^HG6^* (Figure 6b).

**Figure 6.**
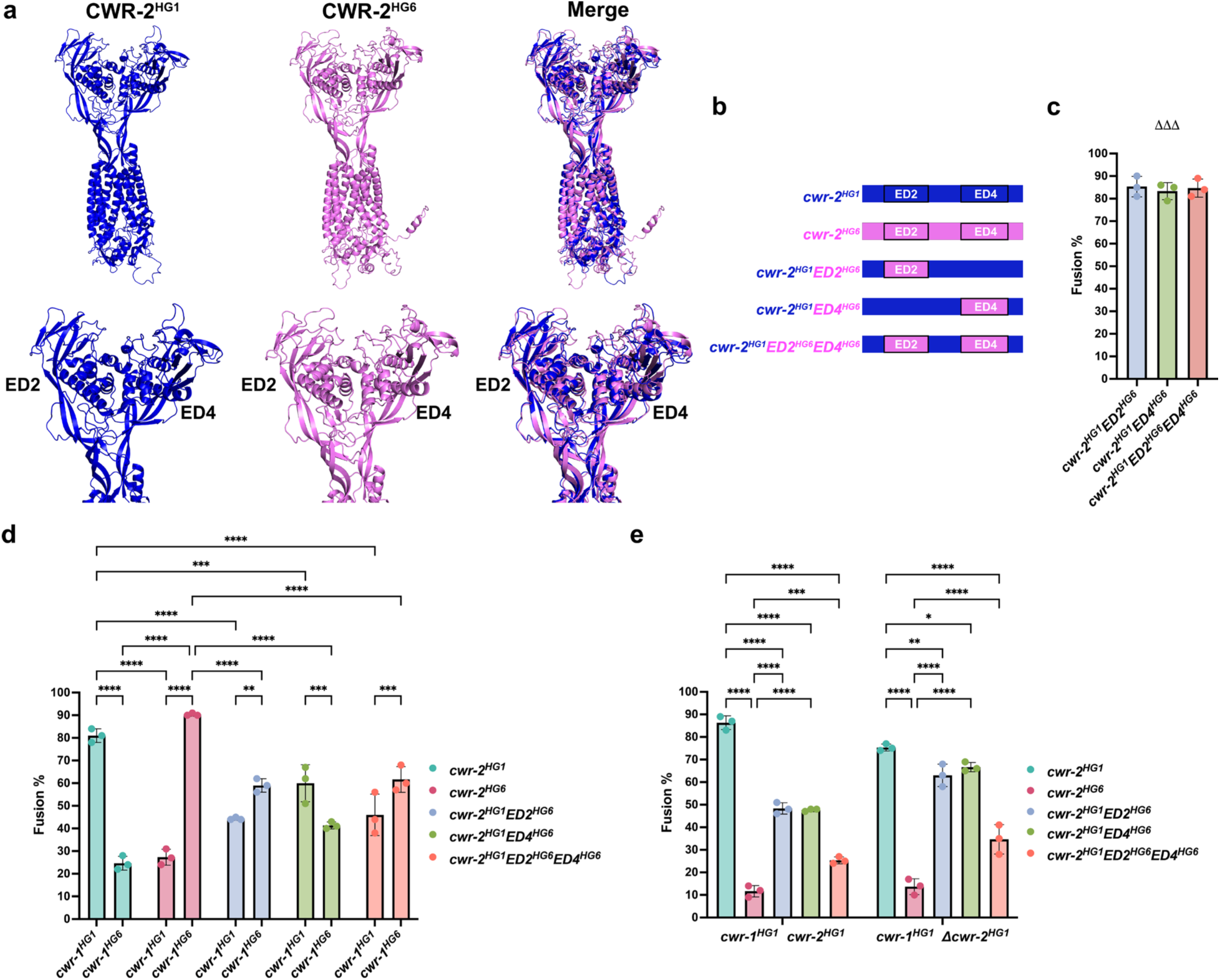
Dissecting the influence of two extracellular domains of CWR-2 in allorecognition at the cell wall remodeling checkpoint. **a**. Predicted three-dimensional structure of CWR-2^HG1^ and CWR-2^HG6^ using ColabFold (Mirdita *et al*. 2022), with a merged image of both structures. The lower panels show a close-up of the CWR-2^HG1^ ED2 and ED4 domains, the CWR-2^HG6^ ED2 and ED4 domains and merged image. **b**. Schematic of CWR-2^HG1^ chimeras bearing ED2^HG6^, ED4^HG6^ or ED2^HG6^/ED4^HG6^ regions. Constructs were targeted to the *his-3* locus. **c**. Cell fusion tests of CWR-2^HG1^ ED2^HG6^, CWR-2^HG1^ ED4^HG6^ and CWR-2^HG1^ ED2^HG6^/ED4^HG6^ chimeric strains paired a strain lacking *cwr-1* and *cwr-2* (*ΔΔΔ*). **d**. Cell fusion tests *cwr-2*^HG1^ ED2^HG6^, *cwr-2*^HG1^ ED4^HG6^ and *cwr-2*^HG1^ ED2^HG6^/ED4^HG6^ chimeric strains and a strain carrying either *cwr-1^HG1^* or *cwr-1^HG6^* alleles. All alleles were targeted to the *his-3* locus. **e**. Cell fusion tests of *cwr-2*^HG1^ ED2^HG6^, *cwr-2*^HG1^ ED4^HG6^ and *cwr-2*^HG1^ ED2^HG6^/ED4^HG6^ chimeric strains paired with *cwr-1^HG1^ cwr-2^HG1^* (FGSC 2489) germlings or with germlings expressing *cwr-1^HG1^*at the native locus (*cwr-1^HG1^ Δcwr-2^HG1^*). A Two-way ANOVA followed by Tukey’s post-hoc test was used for statistical analysis. Error bars represent SD (Standard deviation), *p<0.05, **p<0.01, ***p<0.001, ****p<0.0001, ns: not significant. Cell fusion test experiments were performed in biological triplicate, assessing the fusion of 100 germling pairs for each replicate. Individual p-values are reported in Supplementary Table S4.

All *cwr-2* chimeric strains were first paired with the *ΔΔΔ* mutant; all showed fusion percentages of around 80% (Figure 6c). In contrast, cell fusion percentages in pairings between the *cwr-1^HG1^* + *cwr-2^HG1^ ED2^HG6^*and *cwr-1^HG1^* + *cwr-2^HG1^ ED2^HG6^ED4^HG6^*strains showed fusion percentages of 44% and 46%, respectively (Figure 6d). Fusion percentages between *cwr-1^HG6^* + *cwr-2^HG1^ ED2^HG6^* and *cwr-1^HG6^* + *cwr-2^HG1^ ED2^HG6^ ED4^HG6^* were higher (59% and 62%, respectively). These data suggested that the inclusion of ED2 from HG6 affected allelic specificity of an otherwise CWR-2^HG1^ strain. In contrast, the *cwr-1^HG1^*+ *cwr-2^HG1^ ED4^HG6^* chimeras showed fusion percentages of 60% and 41% in pairings with a *cwr-1^HG6^* strain, indicating that the E4 domain affected fusion, but did not significantly alter *cwr-2* allelic specificity (Figure 6d). These data suggest that the ED2^HG6^ domain of CWR-2 is an important domain for HG6 allorecognition.

The *cwr-2^HG1^* chimeras were also paired with the *cwr-1^HG1^ cwr-2^HG1^* (FGSC 2489) strain as well as a *cwr-1^HG1^ Δcwr-2^HG1^* mutant (*cwr-1* is at the native locus in this strain). In *cwr-1^HG1^ cwr-2^HG1^* + *cwr-2^HG1^ED2^HG6^*and *cwr-1^HG1^ cwr-2^HG1^* + *cwr-2^HG1^ED4^HG6^* pairings, the fusion percentage of ∼48% was obtained (Figure 6e). Pairings where both the ED2 and ED4 regions were swapped (*cwr-1^HG1^ cwr-2^HG1^* + *cwr-2^HG1^ ED2^HG6^ ED4^HG6^*) showed a lower fusion percentage of 25% (Figure 6e). Fusion was higher in pairings with a *cwr-2* deletion strain (*cwr-1^HG1^ Δcwr-2^HG1^* + *cwr-2^HG1^ED2^HG6^*and *cwr-1^HG1^ Δcwr-2^HG1^* + *cwr-2^HG1^ED4^HG6^*) (63% and 67%, respectively), versus pairings between *cwr-1^HG1^ Δcwr-2^HG1^* + *cwr-2^HG1^ ED2^HG6^ ED4^HG6^* (fusion percentages of 35%) (Figure 6e). These data indicated that the *cwr-2^HG1^ ED2^HG6^ ED4^HG6^* chimera functioned more efficiently in switching allelic specificity when paired with *cwr-1^HG1^ cwr-2^HG1^* (FGSC 2489) and cwr*-1^HG6^ Δcwr-2^HG1^*strains, both of which possess the *cwr-1* allele at the native locus.

### Examining the impact of the ED2 and ED4 domains of CWR-2 between two closely related haplogroups (HG3 and HG2) on allorecognition at the cell wall remodeling fusion checkpoint

Data from the chimeric swaps between *cwr-2^HG1^* and *cwr-2^HG6^* indicated that ED2/ED4 extracellular domains impacted *cwr-2* allelic recognition. To test allelic specificity domains in *cwr-2* haplogroups that are closely related, we constructed similar chimeras between *cwr-2^HG2^* and *cwr-2^HG3^* (Supplementary Figure S2). Modeling of the CWR-2^HG2^ and CWR-2^HG3^ by ColabFold (Mirdita *et al*. 2022) is shown in Figure 7a. The RMSD value for this CWR-2 protein pair was 0.950 Å, which is lower than the value of 1.690 Å obtained between HG1 and HG6 CWR-2 proteins. This value indicates that the overall three-dimensional structures of CWR-2 from HG2 and HG3 were quite similar or closely aligned structurally. The RMSD values for the ED2 and ED4 domains were 0.747 Å and 0.895 Å, respectively, slightly higher than those obtained for HG1 and HG6 (0.730 Å and 0.650 Å). The ED2*^HG3^*(G164-R495) and ED4*^HG3^*(F743-A1099) domains were replaced by the corresponding ED2*^HG2^*(G164-R495) and ED4*^HG2^*(F743-A1096) domains from HG2, resulting in the *cwr-2^HG3^ED2^HG2^, cwr-2^HG3^ED4^HG2^,* and *cwr-2^HG3^ED2^HG2^ED4^HG2^*chimeric strains (Figure 7b). All of the *cwr-2* HG3/HG2 chimeras showed identical and high fusion percentages with the *ΔΔΔ* mutant lacking both *cwr-1* and *cwr-2* (Figure 7c). In control pairings, *cwr-1^HG2^* + *cwr-2^HG3^* showed a fusion percentage of 24%, while *cwr-1^HG3^* + *cwr-2^HG2^* pairings showed a fusion percentage of 53% (Figure 7d; Supplementary Figure S1b). These data support our findings that not all pairings of wild type *cwr-1* + *cwr-2* strains in reciprocal combinations have an identical fusion blockage effect. Cell fusion percentages were similar between the control pairing (*cwr-1^HG3^* + *cwr-2^HG2^;* 53%) and *cwr-1^HG3^* + *cwr-2^HG3^* ED2^HG2^, *cwr-1^HG3^* + *cwr-2^HG3^* ED4^HG2^, and *cwr-1^HG3^* + *cwr-2^HG3^* ED2/ED4^HG2^ pairings (43%, 53% and 47% respectively) (Figure 7d). These data showed that the three chimeras exhibited fusion percentages comparable to the control *cwr-1^HG3^* + *cwr-2^HG2^* pairings (53%), indicating that the ED2, ED4 and ED2/ED4 domains play an important role in altering *cwr-2* allelic specificity. In contrast, the *cwr-1^HG2^* + *cwr-2^HG3^* ED2^HG2^, *cwr-1^HG2^* + *cwr-2^HG3^* ED4^HG2^, and *cwr-1^HG2^* + *cwr-2^HG3^* ED2/ED4^HG2^ pairings showed higher fusion percentages (77%, 72%, and 70%, respectively), close to the 80% fusion percentage observed in the control pairing *cwr-1^HG2^* + *cwr-2^HG2^*(Figure 7d). Statistical analysis showed no significant differences in the fusion percentages among the three CWR-2 chimeras. These results support a role for the CWR-2 ED2 and ED4 regions in two different HG pairings (HG1 + HG6 and HG2 + HG3) in determining allelic specificity, even between the closely related HG2 and HG3 strains.

**Figure 7.**
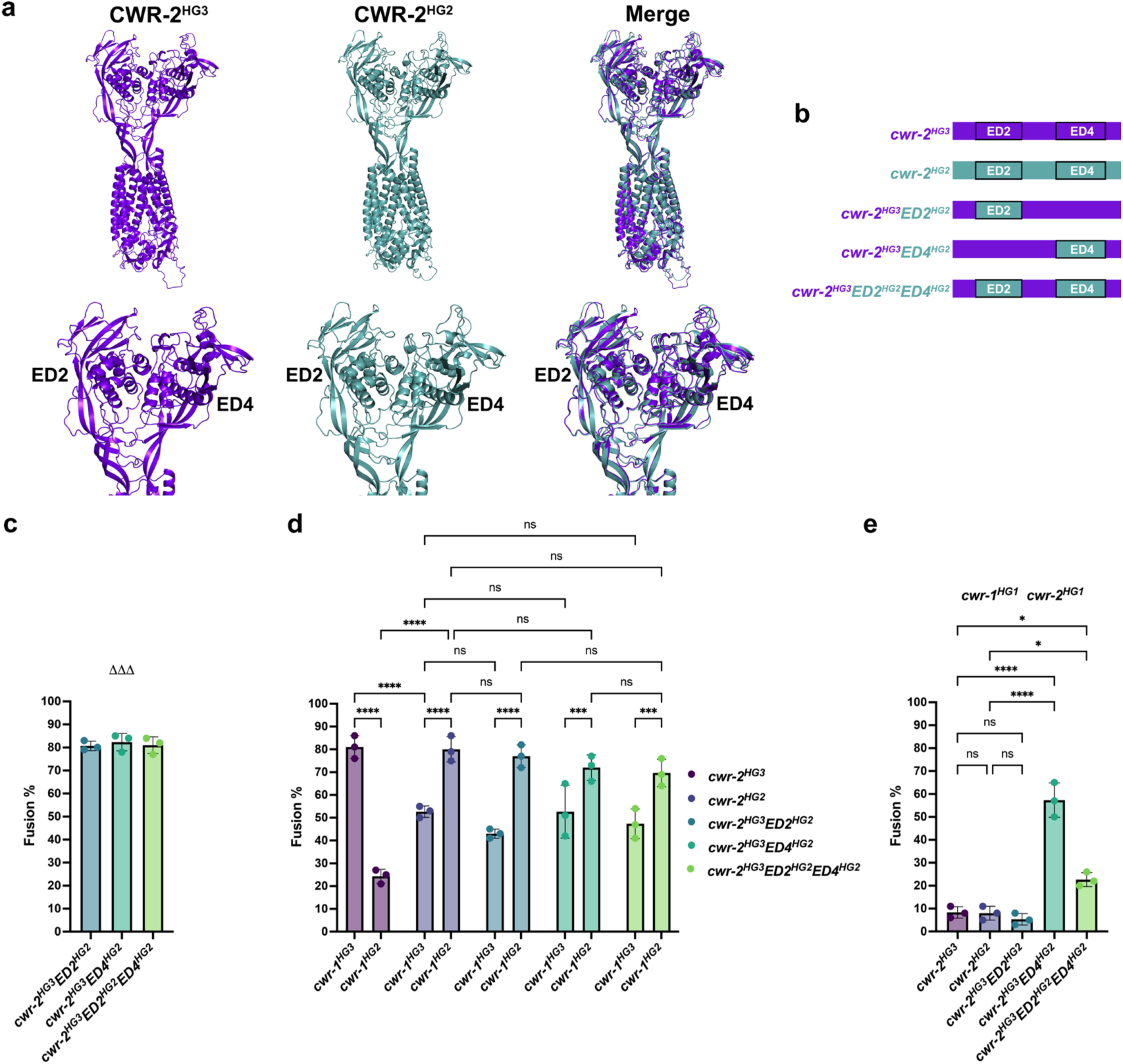
Examining the impact of two large extracellular domains of CWR-2 in allorecognition at the cell wall remodeling checkpoint. **a**. Predicted three-dimensional structure of CWR-2 from HG2 or HG3 using ColabFold (Mirdita *et al*. 2022) and merged image. Lower panels show a close-up of ED2 and ED4 and a merged image emphasizing conformational differences. **b**. Schematic of *cwr-2*^HG3^ ED2^HG2^, ED4^HG2^, or ED2^HG2^/ED4^HG2^ chimeras. Constructs were targeted to the *his-3* locus and under the regulation of the P*tef-1* promoter. **c**. The *cwr-2^HG3^* chimeric strains were paired a Δ*cwr-1* Δ*cwr-2* (ΔΔΔ) mutant. **d**. Analysis of the cell fusion events between the CWR-2^HG3^ chimeric strains and *cwr-1^HG3^* or *cwr-1^HG2^*germlings. A two-way ANOVA followed by Tukey’s post-hoc test was used for statistical analysis. Error bars represent SD (Standard deviation), ***p<0.001, ****p<0.0001, ns: not significant. **e**. Evaluation of the cell fusion events of strains expressing the *cwr-2^HG3^*chimeras paired with *cwr-1^HG1^ cwr-2^HG1^* (FGSC 2489) germlings. As controls, strains expressing *cwr-2^HG3^* or *cwr-2^HG2^* were paired with FGSC 2489. A one-way ANOVA followed by Tukey’s post-hoc test was used for statistical analysis. Error bars represent SD (standard deviation), *p<0.05, ****p<0.0001, ns: not significant. Cell fusion tests were performed in biological triplicate, assessing the fusion events of 100 germling pairs for each replicate. Individual p-values are reported in Supplementary Table S4.

When paired with the wild type strain (*cwr-1^HG1^ cwr-2^HG1^;* FGSC 2489), strains carrying *cwr-2^HG2^* or *cwr-2^HG3^* at the *his-3* locus showed low fusion percentages (8% for both cases; Figure 7e). A pairing between wild type and the *cwr-2^HG3^* ED2^HG2^ chimera (*cwr-1^HG1^ cwr-2^HG1^* + *cwr-2^HG3^* ED2^HG2^) also showed a low fusion percentage (5%). However, the *cwr-1^HG1^ cwr-2^HG1^* + *cwr-2^HG3^*ED4^HG2^ pairing showed a high fusion percentage (57%), while the *cwr-1^HG1^ cwr-2^HG1^* + *cwr-2^HG3^*ED2/ED4^HG2^ pairing showed an intermediate level of fusion (23%) (Figure 7e). These data suggest that the inclusion of the HG2 ED4 domain in an otherwise HG3 CWR-2 protein pushed allelic specificity towards compatibility with an CWR-1 HG1 strain, suggesting that this chimera may represent a novel CWR-2 specificity.

## Discussion

The formation of an interconnected mycelial network via germling/hyphal fusion is a hallmark of filamentous fungi. Formation of this network involves a complex interplay of cell signaling that induces chemotropic interactions that ultimately result in cell fusion (Herzog *et al*. 2015; Fischer and Glass 2019). The chemotropic signal for cell fusion appears to be highly conserved between the distantly related ascomycete fungi (Fleissner *et al*. 2022; Haj Hammadeh *et al*. 2022). Indeed, mutations in genes that disrupt hyphal fusion in *N. crassa* have a similar fusion defect when mutated in other filamentous ascomycete fungi, suggesting that signals and machinery for cell fusion are highly conserved (Scott *et al*. 2018; Goncalves *et al*. 2020). In *N. crassa*, three checkpoints have been identified that affect cell fusion. Two of them, the communication checkpoint and the cell wall remodeling checkpoint, act prior to membrane merger and cytoplasmic mixing (Goncalves and Glass 2020; Goncalves *et al*. 2020). Both of these checkpoints function to negatively regulate the fusion process, as strains with mutations in genes required for these checkpoints undergo chemotropic interactions (communication checkpoint) or cell wall breakdown and membrane merger (cell wall remodeling checkpoint) with strains they were formerly incompatible with (Heller *et al*. 2016; Goncalves *et al*. 2019; Detomasi *et al*. 2022). The loci that regulate the pre- and post-fusion processes in filamentous fungi have hallmarks of being under balancing selection: alleles at these allorecognition loci are highly polymorphic, isolates group into haplogroups, allele frequency in populations are nearly equal and alleles often show trans-species polymorphisms (Klein *et al*. 1998; Goncalves *et al*. 2020). Although a number of allorecognition loci have been identified in filamentous fungi, very few of them have had their molecular recognition mechanisms characterized and regions required for allelic specificity identified; exceptions include the post-fusion death loci, *rcd-1* in *N. crassa*, which encodes a homolog of gasdermin (Daskalov *et al*. 2019; Daskalov *et al*. 2020; Li *et al*. 2024) and *het-s* in *Podospora anserina*, which encodes a prion (Riek and Saupe 2016; Bardin *et al*. 2021; Son 2024).

The most well characterized cell fusion process in fungi is in the yeast species, *Saccharomyces cerevisiae* and *Schizosaccharomyces pombe* (Martin 2019; Clark-Cotton *et al*. 2022; Sieber *et al*. 2023), which occurs during mating. In these species, secretory vesicles carrying cell wall degrading enzymes are localized by the actin cytoskeleton to the point of contact between mating cells (Cappellaro *et al*. 1998; Paterson *et al*. 2008; Huberman and Murray 2014). Regulated secretion of these enzymes is important to prevent premature cell wall digestion, which would result in cell lysis (Philips and Herskowitz 1997; Merlini *et al*. 2013; Clark-Cotton *et al*. 2022). In *N. crassa*, cell wall breakdown and membrane merger are negatively regulated during vegetative cell fusion by CWR-1 and CWR-2, which, in incompatible cells, interact *in trans* to trigger a cell fusion block at the point of cell wall degradation. A key question in this context is how this recognition mechanism occurs. Our initial hypothesis was that CWR-1, which is a chitin polysaccharide monooxygenase, formed HG-specific chitin products in one cell, would be sensed by the transmembrane protein CWR-2 in the plasma membrane of the partner cell. However, this hypothesis was disproved as mutations in CWR-1 that abolish PMO activity were unaffected in the triggering allorecognition and a block in cell fusion (Detomasi *et al*. 2022). We therefore turned our attention to regions of the PMO domain that were conserved within a haplogroup, but divergent between haplogroups and that regulated allelic specificity.

Using ColabFold V1.5.5 AlphaFold2 (Jumper *et al*. 2021; Mirdita *et al*. 2022), the superimposed conformational structures of the PMO domain from CWR-1 from the six different haplogroups highlighted two regions with major structural differences: the LC region and, to a lesser extent, the L2 region. The evaluation of the L2 and LC regions of the PMO domain by using chimeras from two distant HGs (HG1 and HG6) indicated that the L2 and LC regions were essential for conferring allelic specificity (Figures 3, 4; Supplemental Figure S7). Indeed, just the LC region of the PMO, attached to the glycine linker and X278 chitin binding domain was sufficient to alter fusion percentages to a degree (Figure 5), even though there were structural differences that were lacking that are present in an intact PMO domain. The PMO domain regions that affected allelic specificity were reciprocal (i.e. L2 and LC were important for H1 and H6 specificity) further strengthening the support for this region in regulating CWR-1 allorecognition. It is unclear, however, from modeling data of L2 and LC how these regions functions in specificity on a molecular level. However, both the L2 and LC regions reside on the accessible regions of the PMO domain, suggesting that physical interaction, perhaps with CWR-2, may play a role.

The structural prediction of the transmembrane protein CWR-2 helped to clarify the dimensionality of the protein inserted into the membrane and has a much more complex structural conformation than CWR-1. Given the architecture and extension of the two large extracellular domains ED2 and ED4 that were identical in amino acid sequence within a haplogroup, but divergent between haplogroups, we hypothesized that these domains would play a role in conferring selectivity in allorecognition. In distantly related haplogroups (HG1 and HG6), the ED2 domain appeared to have a more prominent role in driving allorecognition shifts. However, when paired with strains carrying the *cwr-1* allele at the native locus (*cwr-1 cwr-2* and *cwr-1 Δcwr-2*), the chimera with both ED2 and ED4 domains replaced exhibited the most pronounced allelic specificity changes. In contrast, for closely related haplogroups (HG2 and HG3), all three chimeras (with replaced ED2, ED4, and ED2/ED4 domains) showed results similar to that of controls (*cwr-1^HG3^ + cwr-2^HG2^* and *cwr-1^HG2^ + cwr-2^HG2^*), suggesting that both domains are important for allelic specificity (Supplementary Figure S7). While our data does not conclusively establish whether ED2 or ED4 contributes to specificity more than the other, the high levels of cellular fusion observed within these closely related haplogroups may be attributed to their inherent similarities despite their haplogroup differences. Interestingly, when these chimeras were paired with the wild-type strain (*cwr-1^HG1^ cwr-2^HG1^*), the chimera *cwr-2^HG3^* ED2^HG2^ displayed the lowest fusion levels, highlighting the importance of the ED2 and aligning with the findings observed in chimeras from the more distant groups (HG1 and HG6).

Evidence from chimeric strains of the CWR-2 indicates that the ED2 and ED4 domains are key determinants of allorecognition specificity. As with the L2 and LC regions of the PMO domain, the ED2 and ED4 were reciprocally important for CWR-2 allelic specificity, although the strength of this function was not as definitive as with CWR-1. Given the structural complexity of the protein, it is likely that other domains also play a role in this process. The alignment among the six different HGs (Supplementary Figure S4) showed that the cytoplasmic domain CD3 exhibits high amino acid variability across the six HGs as compared to other cytoplasmic domains. It is possible that CWR-2 may undergo conformational changes when activated that trigger an internal signal that involve haplogroup specific domains that face the cytoplasm.

Upon contact, cells switch from undergoing chemotropic growth to cell wall breakdown and membrane merger (Herzog *et al*. 2015; Fischer and Glass 2019; Fleissner *et al*. 2022). This process must be carefully regulated to avoid cell lysis during mating and vegetative cell fusion (Jin *et al*. 2004; Fleissner *et al*. 2009; Palma-Guerrero *et al*. 2014; Palma-Guerrero *et al*. 2015). Previously, it was reported that cells that are blocked at the *cwr* checkpoint show increased cell wall deposition and continued signaling of proteins associated with chemotropic growth (Goncalves *et al*. 2019). These observations indicate that the *cwr* checkpoint prevents the switch from chemotrophic growth upon cell contact to cell wall deconstruction and membrane merger. It is unknown, even during yeast mating, how this transition is regulated. The simplest model for function of the *cwr* checkpoint is a physical interaction between CWR-1 and CWR-2 of different HGs that prevents this switch. Previous experiments where wild type compatible germlings were treated with CWR-1 purified from incompatible HGs did not change the cell fusion proficiency of these cells (Detomasi *et al*. 2022), although there are obvious technical aspects of this experiment that were difficult to address. Our work described here on the deciphering the regions of CWR-1 and CWR-2 that regulate specificity provides some insight for further research should concentrate on elucidating the underlying mechanisms of the interactions between CWR-1 and CWR-2 and the signaling pathways involved in regulating cell fusion arrest in incompatible cells.

## Data Availability Statement

All strains generated in this study are available at the Fungal Genetics Stock Center (https://www.plantpath.k-state.edu/research-services/fungal-genetics-stock-center/). Primers used in this study are listed in Supplementary Table S2. The P-values for the cell fusion tests are reported in Supplementary Table S4. RMSD values of the predicted structures of CWR-1 and CWR-2 from the different HGs are reported in Supplementary Table S5.

## Acknowledgements

This work was funded by a United States National Science Foundation Grant (MCB-1818283) to NLG. The authors are grateful to Dr. Tyler Detomasi for his invaluable suggestions for the development of the chimeras and to Maria Mercado for her assistance with lab work.

## References

Afzali, B., G. Lombardi and R. I. Lechler, 2008 Pathways of major histocompatibility complex allorecognition. Curr Opin Organ Transplant 13: 438–444.

Bardin, T., A. Daskalov, S. Barrouilhet, A. Granger-Farbos, B. Salin et al., 2021 Partial prion cross-seeding between fungal and mammalian amyloid signaling motifs. mBio 12.

Cappellaro, C., V. Mrsa and W. Tanner, 1998 New potential cell wall glucanases of *Saccharomyces cerevisiae* and their involvement in mating. J Bacteriol 180: 5030–5037.

Charmetant, X., G. J. Pettigrew and O. Thaunat, 2024 Allorecognition Unveiled: Integrating recent breakthroughs into the current paradigm. Transpl Int 37: 13523.

Chothia, C., and A. M. Lesk, 1986 The relation between the divergence of sequence and structure in proteins. EMBO J 5: 823–826.

Clark-Cotton, M. R., K. C. Jacobs and D. J. Lew, 2022 Chemotropism and cell-cell fusion in fungi. Microbiol Mol Biol Rev 86: e0016521.

Daskalov, A., P. Gladieux, J. Heller and N. L. Glass, 2019 Programmed cell death in *Neurospora crassa* is controlled by the allorecognition determinant *rcd-1*. Genetics 213: 1387–1400.

Daskalov, A., P. S. Mitchell, A. Sandstrom, R. E. Vance and N. L. Glass, 2020 Molecular characterization of a fungal gasdermin-like protein. Proc Natl Acad Sci U S A 117: 18600–18607.

Debets, A. J. M., and A. J. F. Griffiths, 1998 Polymorphism of *het*-genes prevents resource plundering in *Neurospora crassa*. Mycol. Res. 102: 1343–1349.

Debets, F., X. Yang and A. J. Griffiths, 1994 Vegetative incompatibility in Neurospora: its effect on horizontal transfer of mitochondrial plasmids and senescence in natural populations. Curr Genet 26: 113–119.

Detomasi, T. C., A. M. Rico-Ramirez, R. I. Sayler, A. P. Goncalves, M. A. Marletta et al., 2022 A moonlighting function of a chitin polysaccharide monooxygenase, CWR-1, in *Neurospora crassa* allorecognition. Elife 11.

Ellison, C. E., C. Hall, D. Kowbel, J. Welch, R. B. Brem et al., 2011 Population genomics and local adaptation in wild isolates of a model microbial eukaryote. Proc Natl Acad Sci USA 108: 2831–2836.

Fischer, M. S., and N. L. Glass, 2019 Communicate and Fuse: How filamentous fungi establish and maintain an interconnected mycelial network. Front Microbiol 10: 619.

Fleissner, A., S. Diamond and N. L. Glass, 2009 The *Saccharomyces cerevisiae* PRM1 homolog in *Neurospora crassa* is involved in vegetative and sexual cell fusion events but also has postfertilization functions. Genetics 181: 497–510.

Fleissner, A., A. G. Oostlander and L. Well, 2022 Highly conserved, but highly specific: Somatic cell-cell fusion in filamentous fungi. Curr Opin Cell Biol 79: 102140.

Glass, N. L., and I. Kaneko, 2003 Fatal attraction: Nonself recognition and heterokaryon incompatibility in filamentous fungi. Eukaryot. Cell 2: 1–8.

Goncalves, A. P., and N. L. Glass, 2020 Fungal social barriers: to fuse, or not to fuse, that is the question. Commun Integr Biol 13: 39–42.

Goncalves, A. P., J. Heller, A. Daskalov, A. Videira and N. L. Glass, 2017 Regulated forms of cell death in fungi. Front Microbiol 8: 1837.

Goncalves, A. P., J. Heller, A. M. Rico-Ramirez, A. Daskalov, G. Rosenfield et al., 2020 Conflict, competition, and cooperation regulate social interactions in filamentous fungi. Annu Rev Microbiol 74: 693–712.

Goncalves, A. P., J. Heller, E. A. Span, G. Rosenfield, H. P. Do et al., 2019 Allorecognition upon fungal cell-cell contact determines social cooperation and impacts the acquisition of multicellularity. Curr Biol 29: 3006–3017 e3003.

Haj Hammadeh, H., A. Serrano, V. Wernet, N. Stomberg, D. Hellmeier et al., 2022 A dialogue-like cell communication mechanism is conserved in filamentous ascomycete fungi and mediates interspecies interactions. Proc Natl Acad Sci U S A 119: e2112518119.

Heller, J., C. Clave, P. Gladieux, S. J. Saupe and N. L. Glass, 2018 NLR surveillance of essential SEC-9 SNARE proteins induces programmed cell death upon allorecognition in filamentous fungi. Proc Natl Acad Sci U S A 115: E2292–E2301.

Heller, J., J. Zhao, G. Rosenfield, D. J. Kowbel, P. Gladieux et al., 2016 Characterization of greenbeard genes involved in long-distance kind discrimination in a microbial eukaryote. PLoS Biol 14: e1002431.

Hemsworth, G. R., B. Henrissat, G. J. Davies and P. H. Walton, 2014 Discovery and characterization of a new family of lytic polysaccharide monooxygenases. Nat Chem Biol 10: 122–126.

Herzog, S., M. R. Schumann and A. Fleissner, 2015 Cell fusion in *Neurospora crassa*. Curr Opin Microbiol 28: 53–59.

Hickey, P. C., D. Jacobson, N. D. Read and N. L. Louise Glass, 2002 Live-cell imaging of vegetative hyphal fusion in *Neurospora crassa*. Fungal Genet Biol 37: 109–119.

Hickey, P. C., S. R. Swift, M. G. Roca and N. D. Read, 2004 Live-cell imaging of filamentous fungi using vital fluorescent dyes and confocal microscopy., pp. 63–87 in Methods in Microbiology, edited by T. Savidge and C. Pothoulakis. Elsevier, London, U.K.

Huberman, L. B., and A. W. Murray, 2014 A model for cell wall dissolution in mating yeast cells: polarized secretion and restricted diffusion of cell wall remodeling enzymes induces local dissolution. PLoS One 9: e109780.

Huene, A. L., S. M. Sanders, Z. Ma, A. D. Nguyen, S. Koren et al., 2022 A family of unusual immunoglobulin superfamily genes in an invertebrate histocompatibility complex. Proc Natl Acad Sci U S A 119: e2207374119.

Jin, H., C. Carlile, S. Nolan and E. Grote, 2004 Prm1 prevents contact-dependent lysis of yeast mating pairs. Eukaryotic Cell 3: 1664–1673.

Jumper, J., R. Evans, A. Pritzel, T. Green, M. Figurnov et al., 2021 Highly accurate protein structure prediction with AlphaFold. Nature 596: 583–589.

Kabsch, W., 1976 A solution for the best rotation to relate two sets of vectors. Acta Crystallographica Section A 32: 922–923.

Karadge, U. B., M. Gosto and M. L. Nicotra, 2015 Allorecognition proteins in an invertebrate exhibit homophilic interactions. Curr Biol 25: 2845–2850.

Katoh-Kurasawa, M., P. Lehmann and G. Shaulsky, 2024 The greenbeard gene tgrB1 regulates altruism and cheating in *Dictyostelium discoideum*. Nat Commun 15: 3984.

Kessin, R. H., 2001 Dictyostelium: Evolution, Cell Biology and the Development of Multicellularity. Cambridge University Press, Cambridge

Klein, J., A. Sato, S. Nagl and C. O’hUigin, 1998 Molecular trans-species polymorphism. Annu. Rev. Ecol. Syst. 29: 1–21.

Kufareva, I., and R. Abagyan, 2012 Methods of protein structure comparison. Methods Mol Biol 857: 231–257.

Li, Y., Y. Hou, Q. Sun, H. Zeng, F. Meng et al., 2024 Cleavage-independent activation of ancient eukaryotic gasdermins and structural mechanisms. Science 384: adm9190.

Margolin, B. S., M. Freitag, and E.U. Selker 1997 Improved plasmids for gene targeting at the *his-3* locus of *Neurospora crassa* by electroporation. Fungal Genetics Newsletter 34: 34–36

Marino, J., J. Paster and G. Benichou, 2016 Allorecognition by T lymphocytes and allograft rejection. Front Immunol 7: 582.

Martin, S. G., 2019 Molecular mechanisms of chemotropism and cell fusion in unicellular fungi. J Cell Sci 132.

Merlini, L., O. Dudin and S. G. Martin, 2013 Mate and fuse: how yeast cells do it. Open Biol 3: 130008.

Mirdita, M., K. Schutze, Y. Moriwaki, L. Heo, S. Ovchinnikov et al., 2022 ColabFold: making protein folding accessible to all. Nat Methods 19: 679–682.

Palma-Guerrero, J., C. R. Hall, D. Kowbel, J. Welch, J. W. Taylor et al., 2013 Genome wide association identifies novel loci involved in fungal communication. PLoS Genet 9: e1003669.

Palma-Guerrero, J., A. C. Leeder, J. Welch and N. L. Glass, 2014 Identification and characterization of LFD1, a novel protein involved in membrane merger during cell fusion in *Neurospora crassa*. Mol Microbiol 92: 164–182.

Palma-Guerrero, J., J. Zhao, A. P. Goncalves, T. L. Starr and N. L. Glass, 2015 Identification and characterization of LFD-2, a predicted fringe protein required for membrane integrity during cell fusion in *Neurospora crassa*. Eukaryot Cell 14: 265–277.

Paterson, J. M., C. A. Ydenberg and M. D. Rose, 2008 Dynamic localization of yeast Fus2p to an expanding ring at the cell fusion junction during mating. J Cell Biol 181: 697–709.

Philips, J., and I. Herskowitz, 1997 Osmotic balance regulates cell fusion during mating in *Saccharomyces cerevisiae*. J Cell Biol 138: 961–974.

Phillips, C. M., W. T. Beeson, J. H. Cate and M. A. Marletta, 2011 Cellobiose dehydrogenase and a copper-dependent polysaccharide monooxygenase potentiate cellulose degradation by *Neurospora crassa*. ACS Chem Biol 6: 1399–1406.

Rico-Ramirez, A. M., A. P. Goncalves and N. Louise Glass, 2022 Fungal cell death: The beginning of the end. Fungal Genet Biol 159: 103671.

Riek, R., and S. J. Saupe, 2016 The HET-S/s prion motif in the control of programmed cell death. Cold Spring Harb Perspect Biol 8.

Rodriguez-Valbuena, H., J. Salcedo, O. De Their, J. F. Flot, S. Tiozzo et al., 2024 Genetic and functional diversity of allorecognition receptors in the urochordate, *Botryllus schlosseri*. BioRxiv.

Rosengarten, R. D., and M. L. Nicotra, 2011 Model systems of invertebrate allorecognition. Curr Biol 21: R82–92.

Saupe, S. J., 2000 Molecular genetics of heterokaryon incompatibility in filamentous ascomycetes. Microbiol Mol Biol Rev 64: 489–502.

Schindelin, J., I. Arganda-Carreras, E. Frise, V. Kaynig, M. Longair et al., 2012 Fiji: an open-source platform for biological-image analysis. Nat Methods 9: 676–682.

Scott, B., K. Green and D. Berry, 2018 The fine balance between mutualism and antagonism in the Epichloe festucae-grass symbiotic interaction. Curr Opin Plant Biol 44: 32–38.

Sieber, B., J. M. Coronas-Serna and S. G. Martin, 2023 A focus on yeast mating: From pheromone signaling to cell-cell fusion. Semin Cell Dev Biol 133: 83–95.

Son, M., 2024 A story between s and S: [Het-s] prion of the fungus *Podospora anserina*. Mycobiology 52: 85–91.

van Diepeningen, A. D., A. J. Debets and R. F. Hoekstra, 1997 Heterokaryon incompatibility blocks virus transfer among natural isolates of black Aspergilli. Curr Genet 32: 209–217.

Vogel, H. J., 1956 A convenient growth medium for *Neurospora*. Microbiol. Genet. Bull. 13: 42–46.

Westergaard, M., and H. K. Mitchell, 1947 Neurospora V. A synthetic medium favoring sexual reproduction. Amer. J. Bot. 34: 573–577.

